# Age-dependent glycomic response to the 2009 pandemic H1N1 influenza virus and its association with disease severity

**DOI:** 10.1101/2020.06.22.165613

**Authors:** Shuhui Chen, Brian Kasper, Bin Zhang, Lauren P. Lashua, Ted M. Ross, Elodie Ghedin, Lara K. Mahal

## Abstract

Influenza A viruses cause a spectrum of responses, from mild cold-like symptoms to severe respiratory illness and death. Viral strains and intrinsic host factors, such as age, can influence the severity of the disease. Glycosylation plays a critical role in influenza pathogenesis, however the molecular drivers of influenza outcomes remain unknown. In this work, we characterized the glycomic response to the H1N1 2009 pandemic influenza A virus in age-dependent severity. Using a ferret model and a lectin microarray technology we have developed, we compared responses in newly weaned and aged animals, a model for young children and the elderly, respectively. Glycomic analysis revealed changes in glycosylation over the course of the infection, that were associated with severity in an age-dependent manner. These responses may help explain the differential susceptibility to influenza A virus infection of young children and the elderly.

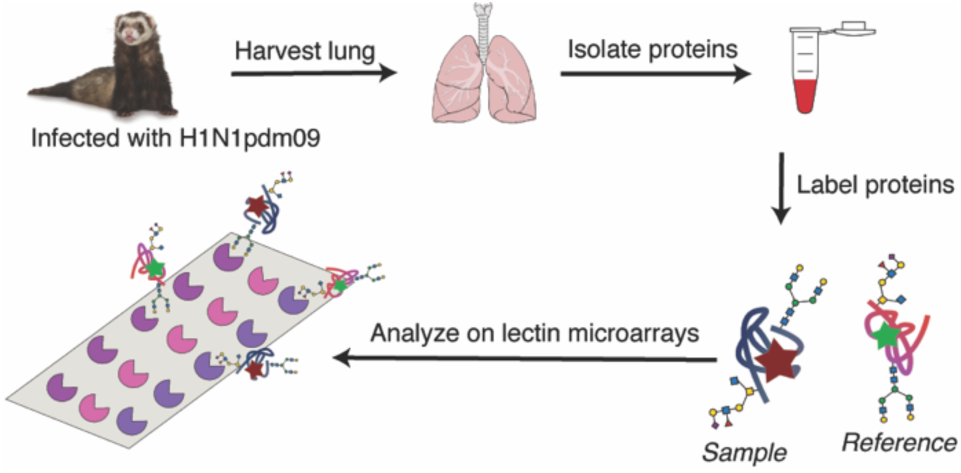

## INTRODUCTION

Influenza A viruses cause a spectrum of responses from mild cold-like symptoms to severe respiratory illness and death.^1^ Both viral strains and intrinsic host factors, such as age, influence disease severity.^2^ In general, infants, young children and individuals older than 65 are more likely to have severe clinical outcomes upon influenza infection. However, during pandemic outbreaks, influenza viral strains can emerge that shift the age-dependent infection patterns. In 2009, a novel H1N1 influenza strain (H1N1pdm09) emerged that caused severe clinical symptoms more frequently in older children and adults than in young children and the elderly.^3^ The lack of response in the elderly was attributed to pre-immunity due to prior waves of seasonal and pandemic influenza strains generating neutralizing antibodies.^4-7^

The ferret model of influenza mimics clinical outcomes for the H1N1pdm09 across the age spectrum.^8-10^ Ferrets are an attractive animal model for influenza as they share similar lung physiology to humans and recapitulate the pathology observed in clinical populations. In addition, human influenza viruses efficiently replicate in ferrets without the need for adaptation, enabling study of the original viral strains.^11^ This is due to the fact that ferrets contain the receptor for human influenza, α-2,6-sialic acids, in their upper respiratory tracts. The similarities between the glycans in the respiratory tract of ferrets to those observed in humans also make this a good model to study glycomic changes in response to influenza infection.^12, 13^

Glycosylation plays an important role in influenza infection and host response.^14, 15^ Adhesion of viral hemagglutinin (HA) to host receptors containing terminal α-2,6-sialic acids located in the upper respiratory tract is the first step in the infection process. Release of viral particles requires viral neuraminidases (NA) to cleave sialic acids.^16^ This enzyme is the target of current therapeutics against influenza.^17, 18^ Due to the importance of HA, NA and sialic acid in influenza biology, the majority of glycomic analysis has focused on these molecules. In previous work, our laboratory used lectin microarray technology to systematically examine the glycomic response to H1N1pdm09 in an adult ferret population.^19^ We found that severity was not related to sialic acid levels, but rather was correlated with the expression of high mannose. This glycan motif is the recognition element for a host of innate lectins that activate the immune system. The complement inducing lectin MBL2, which binds this induced glycan epitope has been strongly associated with influenza severity.^20, 21^ We hypothesize that the induced high mannose may play a causative role in the damage observed from MBL2 levels in influenza.

In this work, we profile glycomic changes, including high mannose, over the course of infection with the pandemic 2009 H1N1 strain of influenza and determine whether they overlay differences in severity observed with age in the ferret model. Since their introduction in 2005,^22^ lectin microarrays have been used to perform glycomic analysis on a wide variety of samples from microvesicles^23^ to human tissues.^24^ Herein, we analyze the glycomic response to H1N1pdm09 infection in newly weaned and aged ferrets using our dual-color lectin microarray technology.^25^ We also compare the baseline glycomes of newly weaned, adult, and aged ferrets. Our data suggest that differences in the glycome, in particular in the high mannose response, may help explain the variation in clinical outcomes as a result of influenza infection.

## METHODS

### Infection of ferrets with H1N1pdm09

Fitch ferrets (*Mustela putorius furo*, female) were obtained from Triple F Farms (Sayre, PA) and verified as negative for antibodies to circulating influenza A (H1N1 and H3N2) and B viruses. Newly weaned ferrets were defined as 6-7 weeks of age, adult ferrets as 6-12 months of age, and aged ferrets as 5.5-7 years of age. Ferrets were pair-housed in stainless steel cages (Shor-Line, KS) containing Sani-Chips laboratory animal bedding (P.J. Murphy Forest Products, NJ) and provided with food and fresh water *ad libitum*. Newly weaned and adult ferrets were administered intranasally the H1N1pdm09 virus, A/California/07/2009 (Influenza Reagents Resources (IRR), BEI Resources, the Centers for Disease Control and Prevention, VA) at a dose of 10^6^ PFU. Aged ferrets were infected at a dose of 10^5^ PFU. The animals were monitored daily for weight loss and disease symptoms, including elevated temperature, low activity level, sneezing, and nasal discharge. Animals were randomly assigned to be sacrificed at day 1, 3, 5, 8, or 14 post-infection (DPI) or euthanized if their clinical condition (e.g., loss of >20% body weight) required it.

### Tissue Collection

Necropsies were performed to collect lung tissue. Lungs were rinsed with cold PBS and the right upper and lower lung lobes were removed. Each lobe was sectioned into quadrants and snap frozen prior to preparation for lectin microarray analysis.

### Definition of Severity Metrics

Severity of infection was determined for all animals studied who were sacrificed at day 8 post-infection. We analyzed two different metrics: weight loss and pneumonia composite score (PCS). In both cases, we determined the cutoffs for mild, moderate and severe by quartile wherein mild = Q0-Q1, moderate = Q1-Q3, severe = Q3-Q4. Weight loss cutoffs for adult ferrets: mild (weight loss <10.8%), moderate (weight loss: 10.8%-16.5%) and severe (weight loss >16.5%). Cutoffs for aged ferrets: mild (weight loss <18.3%), moderate (weight loss: 18.3%-21.9%) and severe (weight loss > 21.9%). PCS cutoffs newly weaned ferrets: mild (PCS < 8.5), moderate (PCS: 8.5-10.5) and severe (PCS > 10.5). For adult ferrets: mild (PCS <8), moderate (PCS: 8-10) and severe (PCS > 10). For aged ferrets: mild(PCS < 8.5), moderate (PCS: 8.5-11), and severe (PCS > 11).

### Lectin microarray

Ferret lung tissue samples were washed with PBS supplemented with protease inhibitors cocktails (PIC) and sonicated on ice in PBS with PIC until completely homogenous. The homogenized samples were then labeled with Alexa Fluor 555-NHS as previously described.^26^ Reference sample was prepared by mixing equal amounts (by total protein) of all samples and labeling with Alexa Fluor 647-NHS (Thermo Fisher). **Supplementary Table 1** summarizes the print list for each experiment. Printing, hybridization and data analysis were performed as previously described.^27^

### Data Availability

Data is available at DOI: 10.7303/syn22176606.

## RESULTS

### Ferret lung glycosylation alters with age

Clinical outcomes and susceptibility to influenza infection vary with age.^28, 29^ Given the role host glycans play in influenza infection and immune response, we characterized the baseline glycosylation in the lungs of uninfected female ferrets from three age categories: newly weaned, adult and aged. Newly weaned ferrets were defined as those from 6-7 weeks of age. The course of H1N1pdm09 infection in this ferret population mirrors that observed clinically in young children, i.e. a mild to moderate presentation of symptoms.^9^ Adult ferrets, defined as 6-12 months of age, have more varied responses to H1N1pdm09 infection ranging from mild to severe, again in line with human populations.^8^ Aged ferrets, defined as 5.5-7 years of age, presented with severe outcomes in >50% of the population when infected with H1N1pdm09.^10^ This is not in line with what was observed clinically in the aged human population for this influenza strain, however, this difference is accounted for by pre-existing immunity to an ancestor of the H1N1pdm09 strain.^4, 6^ In general, aged populations are known to have more severe outcomes upon infection with influenza A viruses.^30, 31^

We analyzed the baseline expression level of host glycans prior to influenza virus infection in lung punch biopsies from newly weaned (NW, n=6), adult (AD, n=4) and aged ferrets (AG, n=3) using our dual-color lectin microarray technology (**Scheme 1**).^25, 26^ Biopsies from both the upper and lower lobes were analyzed for each ferret. Lectin microarrays utilize carbohydrate-binding proteins with well-defined specificities to detect glycan changes between samples.^22, 25-27^ In brief, frozen tissue samples were sonicated and labeled with either Alexa Fluor 555-NHS (sample), or Alexa Fluor 647-NHS (reference). A pooled reference consisting of all samples was used. Equal amounts of sample and reference were analyzed on the lectin microarrays (>100 probes, **Supplementary Table 1**). A heatmap of the normalized data, ordered by ferret age, is shown in **Figure 1a**.

**Figure 1.**
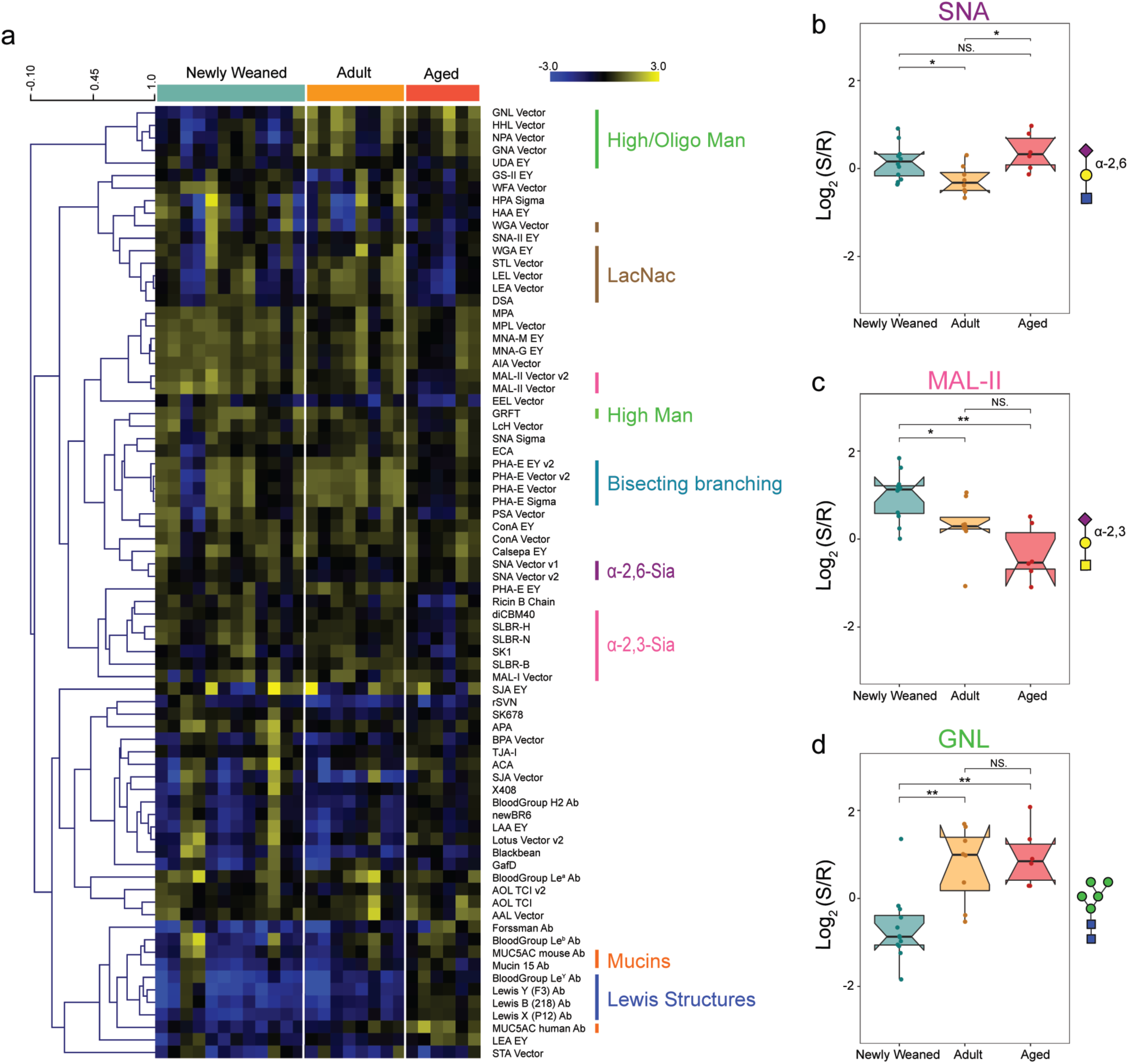
Baseline comparisons between different age groups of uninfected ferrets. a) Heat map of lectin microarray data. Median normalized log_2_ ratios (Sample (S)/Reference(R)) of ferret lung samples were ordered by age (Newly weaned, n = 6, 2 samples per ferret; Adult, n = 4, 2 samples per ferret; Aged, n = 3, 2 samples per ferret). Yellow, log_2_(S) > log_2_(R); blue, log_2_(R) > log_2_(S). Lectins binding α-2,3-sialosides (pink), α-2,6-sialosides (purple), high/oligo-mannose (green), bisecting branching (turquoise), N-Acetyl-D-Lactosamine (brown), mucins (orange) and Lewis structures (blue) are highlighted to the right of the heatmap. b) Boxplot analysis of lectin binding by SNA (α-2,6-sialosides). c) Boxplot analysis of lectin binding by MAL-II (α-2,3- sialosides). d) Boxplot analysis of lectin binding by GNL (oligo-mannose) as a function of age. Newly weaned: cyan; Adult: orange; Aged: red. NS.: Not statistical; *: *p*<0.05; **: *p*<0.01; ***: *p*<0.001. Wilcoxon’s t-test. Glycans bound by lectins are shown in the Symbolic Nomenclature for Glycomics (SNFG) at the side of the boxplots.

Sialic acid glycoconjugates on the host cell surface play a crucial role as both the receptor for influenza binding by hemagglutinin (*α*-2,6-sialosides) and as the target of the influenza neuraminidase (*α*-2,3- and *α*-2,6 sialosides). Differences in sialoside levels may play a role in differential susceptibility to the virus with age. In light of this we examined the expression level of α-2,3- and α-2,6-sialosides in the ferret lung. Both newly weaned and aged ferret lungs had higher levels of α-2,6-sialosides than adult lungs (lectins: SNA, TJA-I, ∼ 1.3-fold NW:AD, ∼1.4- fold AG:AD, **Figure 1b** and **Supplementary Figure S1**), indicating that they may be more susceptible to viral infection. In contrast to α-2,6-sialic acid, we observed a significant age-dependent decrease in α-2,3-sialosides (lectin: MAL-II (∼1.5-fold NW:AD, ∼2-fold NW:AG), SLBR-N,^32^ diCBM40, SLBR-H,^32^ **Figure 1c** and **Supplementary Figure S2**). Of particular note, the lectin MAL-II mainly recognizes α-2,3-sialic acids on *O*-linked glycans. These glycans commonly decorate proteins such as mucins, whose heavily sialylated structures often act as a trap for viral particles.^33^ While a decrease in α-2,3-sialosides is observed with age, we also observe an increase in mucin levels in the lungs (Antibodies: MUC5AC, MUC15, **Supplementary Figure S3**). In addition, we observe a potentially related increase in Lewis structures (Le), which are often observed on mucins (Antibodies: Le^b^, Le^X^, Le^Y^, **Supplementary Figure S4**). Taken together, this suggests that the protective, highly sialylated mucins,^34-36^ that allow clearance of the virus from the lungs, may alter with age in a way that diminishes their protective capacity. This data correlates well with the differences in viral clearance observed in these ferrets, with young ferrets showing more rapid clearance and aged ferrets showing prolonged infection and lower clearance of the virus upon infection.^10^

Recently our group identified high mannose as a severity marker for influenza H1N1pdm09 infection in adult ferrets.^19^ High mannose, defined here as Man_7-9_, and oligo-mannose (defined as Man_5-7_) are targets for innate immune lectin binding and may play a direct role in the damage observed in influenza. Analysis of the mannose binding lectins on the array showed distinct profiles for levels of high- and oligo-mannose.^37^ Uninfected newly weaned animals had significantly lower levels of oligo-mannose (GNL, NPA, HHL, **Figure 1d** and **Supplementary Figure S5**) than either adult or aged ferrets. However, high-mannose levels, recognized by the anti-viral lectin Griffithsin (GRFT),^38^ were higher in the newly weaned and adult animals as compared to the aged ferrets. Whether age-dependent differences in the baseline levels of mannose in uninfected animals have any bearing on the levels of the host response to influenza is currently unclear.

### Aged ferrets have a distinct glycomic response from adults to influenza infection

Disease progression in adult and aged ferrets infected with H1N1pdm09 differ dramatically in timing and presentation.^10^ Aged ferrets had more persistent viral infections and were unable to clear the virus by day 8 post-infection, in contrast to adults. They also developed pneumonia later in the course of their illness and had higher rates of mortality. To study the impact of glycosylation on the host response to influenza in the aged population, we performed lectin microarray analysis on lung punch biopsies from infected aged ferrets (age > 5.5 years). We performed two concurrent studies, a severity study at day 8 post-infection (dpi 8, n = 29 ferrets) and a time-course analysis that included the day 8 samples (dpi 1: n = 5; dpi 3: n = 7; dpi 5: n = 12 ferrets, **Scheme 2**). Uninfected aged ferrets were used as a control (n=3 ferrets). We analyzed 2 samples for each ferret. The animals analyzed in this study correspond to the animals for which pathology data was obtained in Bissel et al.^10^ In our previous work, weight loss was used to evaluate the severity of illness.^10, 19^ This is a common metric for severity in animal models and can only be used for timepoints > day 5 post-infection. In studying the disease progression in aged ferrets we observed a disparity between the pathology and the weight loss data. In this age group, a severe outcome by weight loss was defined as >20% loss of mass.^10^ A pneumonia composite score (PCS), which sums the scores for lung involvement, bronchial severity, and alveolar severity was used to evaluate severity by pathology.^10^ In aged animals, which present with pneumonia at later stages of infection, degree of severity by pathology cannot be determined for early timepoints. A comparison of the severity as defined by weight loss quartiles (Q0-Q1:<18.3%, Q1-Q3: 18.3%-21.9%, Q3-Q4: >21.9%, **Supplementary Figure S6**) to the pneumonia composite scores at day 8 showed no concordance between the two metrics. Instead, in the aged population the trend was towards lower weight loss with increased severity by pathology. This is in direct opposition to the trend observed in adult ferrets. Based on our analysis, weight loss was not an appropriate metric for severity in aged ferrets. Thus, the pneumonia composite score was used as the severity metric for this cohort, as it represents the physiologically relevant degree of illness in influenza infection. We used quartiles to define mild, moderate, and severe infections in the aged population (Mild: PCS < 8.5, n=6 ferrets; Moderate: PCS = 8.5-11, n=15; Severe: PCS > 11, n=8). A heatmap of the normalized data for ferrets sacrificed at day 7-8 post-infection arranged by severity is presented in **Figure 2**. A heatmap of the time-course analysis for aged ferrets is shown in **Supplementary Figure S7**.

**Figure 2.**
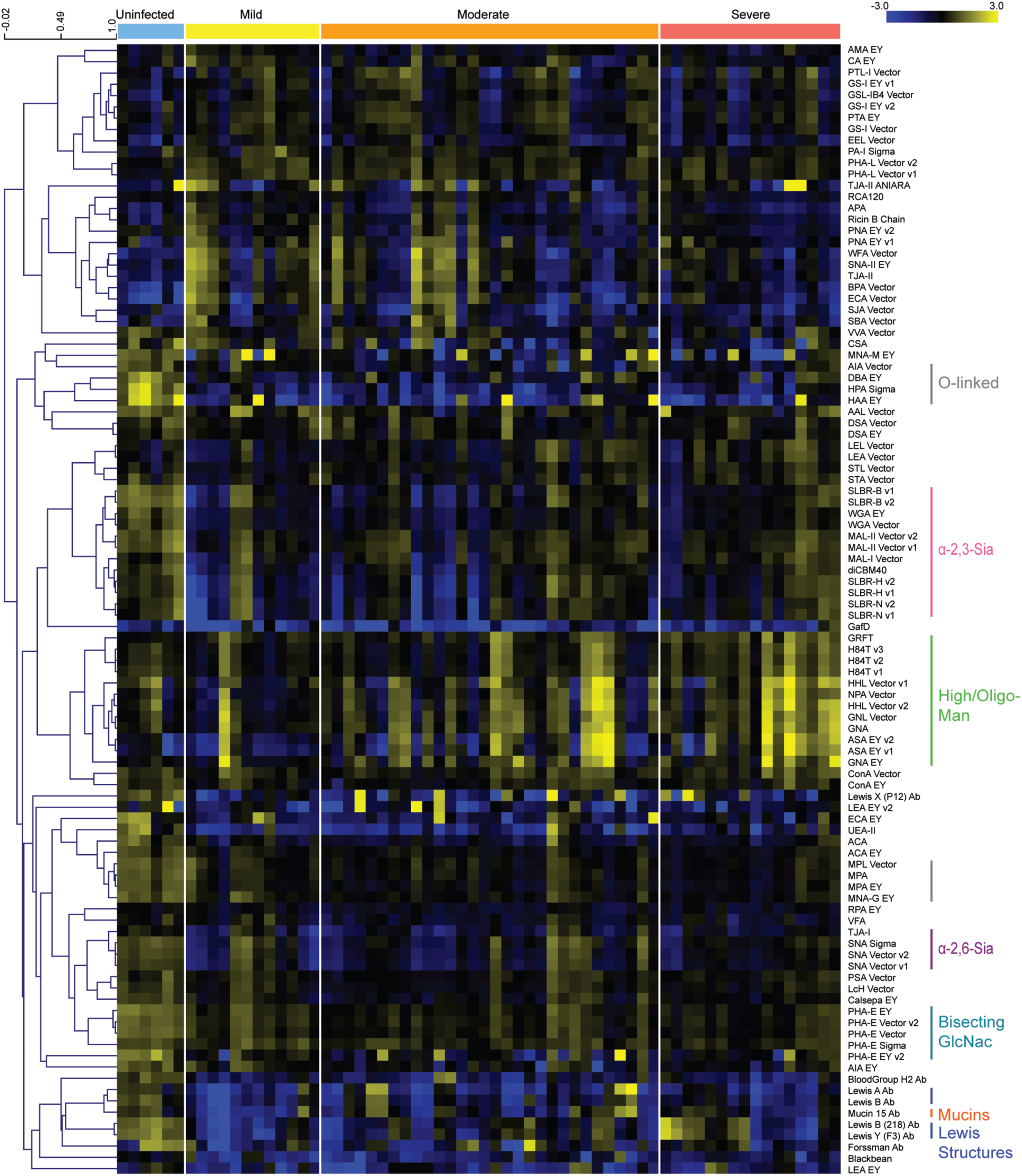
Glycomic changes in aged ferrets in response to influenza infection. Heat map of lectin microarray data for aged ferrets (>5.5 years) infected with H1N1pdm09. Median normalized log_2_ ratios (Sample (S)/Reference(R)) of ferret lung samples were ordered by severity. Uninfected (blue): n = 3. DPI 8: Mild (yellow), n = 6; Moderate (orange), n = 15; Severe (red), n = 8. 2 samples per ferret. Yellow, log_2_(S) > log_2_(R); Blue, log_2_(R) > log_2_(S). Lectins binding α-2,3-sialosides (pink), α-2,6-sialosides (purple), high/oligo-mannose (green), bisecting GlcNac (turquoise), *O*- linked glycans (charcoal) and Lewis structures (blue) are highlighted to the right of the heatmap.

We observed several changes in aged ferrets as a function of infection that did not overlay onto severity. For ease of comparison between the aged and adult host-responses, we reanalyzed the lectin microarray data from our adult ferret cohort^19^ using PCS as the severity metric. In our previous work, adult ferrets showed an increase in α-2,6 sialylation at early infection timepoints, with levels returning to baseline by day 8 post-infection. In contrast, we see a significant loss of α-2,6 sialylation in the aged ferret lungs with this loss occurring early in the course of infection (SNA, TJA-I, **Figure 2** and **Supplementary Figure S8**). We also observe a loss in *α*-2,3-sialosides by day 8 post-infection in aged animals (SLBR-B, SLBR-H, SLBR-N, diCBM-40, MAL-I, MAL-II, **Figure 2** and **Supplementary Figure S9**). A similar loss of *α*-2,3-sialosides was observed in adult ferrets. However, in this population, the response was immediate, while in the aged animals a more gradual loss of this glycan was observed. Influenza neuraminidases are known to prefer *α*- 2,3-sialosides,^39, 40^ and the more gradual loss of this glycan in aged ferrets maps onto the pattern of delayed infection observed in the lungs.^10^

Uninfected aged animals had higher levels of non-sialylated mucins compared to adults. Upon infection, we observed a substantial loss of *O*-linked glycans, Lewis structures, and mucins in aged ferrets (AIA, HAA, HPA, MPA, MNA-G, Mucin 15, Le^a^, Le^b^, Le^Y^, **Figure 2** and **Supplementary Figure S10**). This is similar to previous observations made in adult ferrets, however, the degree of host response in the aged ferrets was more pronounced. We observed a strong loss of *O*-linked glycans, Lewis structures and mucins at the earliest time points (**Supplementary Figure S11**), which may contribute to the inability of older animals to effectively fight infection.

Our previous work suggested a role for high/oligo-mannose as a key mediator of influenza severity.^19^ In our original analysis, severity in the adult ferrets was defined by weight loss. Re- analysis of this data using pneumonia composite scores did not change the previously observed relationship between high/oligo-mannose levels and severity **(Supplementary Figure S12)**. We observe a similar increase in high mannose as a function of severity in the aged animals when severity is defined by pathology (GRFT, HHL, H84T,^41^ NPA, GNA, ASA, **Figures 2** and **Figure 3** and **Supplementary Figure S12**). However, the timing of high mannose induction in the aged ferrets is delayed when compared to adults. In the adult ferrets, high mannose was induced at day 1 post-infection in line with the damage observed by pathology.^10^ In the aged ferrets, we observe significant changes in high mannose by day 3, which persisted to day 8 post-infection. This is consistent with the more delayed development of pneumonia in these animals. Overall, this data supports high mannose as a marker and potential mediator of severity in the host response to influenza infection.

**Figure 3.**
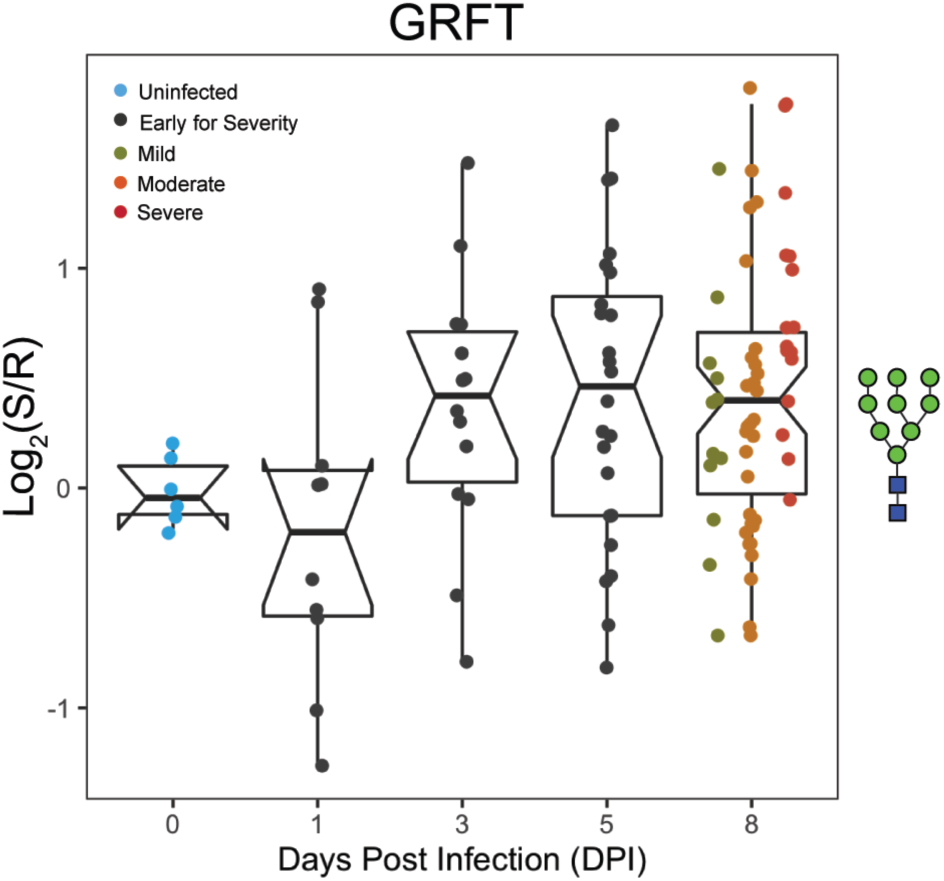
Dynamic changes in high-mannose are observed upon infection in aged ferrets. Time-course analysis of high-mannose as observed by GRFT following H1N1pdm09 infection (t = 0, 1, 3, 5 and 8 days). Boxplot of median normalized log_2_ ratios (Sample (S)/Reference(R)) is shown. Disease severity is indicated by color (Uninfected: blue; Early for Severity: black; Mild: dark yellow; Moderate: orange, Severe: red). Glycans bound by GRFT are shown in the Symbolic Nomenclature for Glycomics (SNFG) at the side of the boxplots.

### Newly weaned ferrets show no severity-dependent changes in the glycome

The 2019 pandemic H1N1 virus presented with mild clinical symptoms in young children.^42-44^ H1N1pdm09 infection in newly weaned ferrets mimic these clinical symptoms, with lower fevers and less weight loss.^9^ Newly weaned ferrets clear virus faster and have much lower levels of alveolar pneumonia.^10^ We analyzed the glycosylation changes in the lungs of newly weaned ferrets (6-7 weeks) infected with H1N1pdm09 for which we had previously obtained pathology data.^10^ To examine the changes in glycosylation over the course of the infection, we performed a time-course study (uninfected, n = 16; dpi 3, n = 4; dpi 5, n = 4; dpi 8, n = 42; dpi 14, n = 3, 2 samples per ferret). Severity was determined for day 8 samples using the PCS metric as previously described. A heatmap of the time-course data, organized by severity for day 8 post-infection, is shown in **Figure 4**.

**Figure 4.**
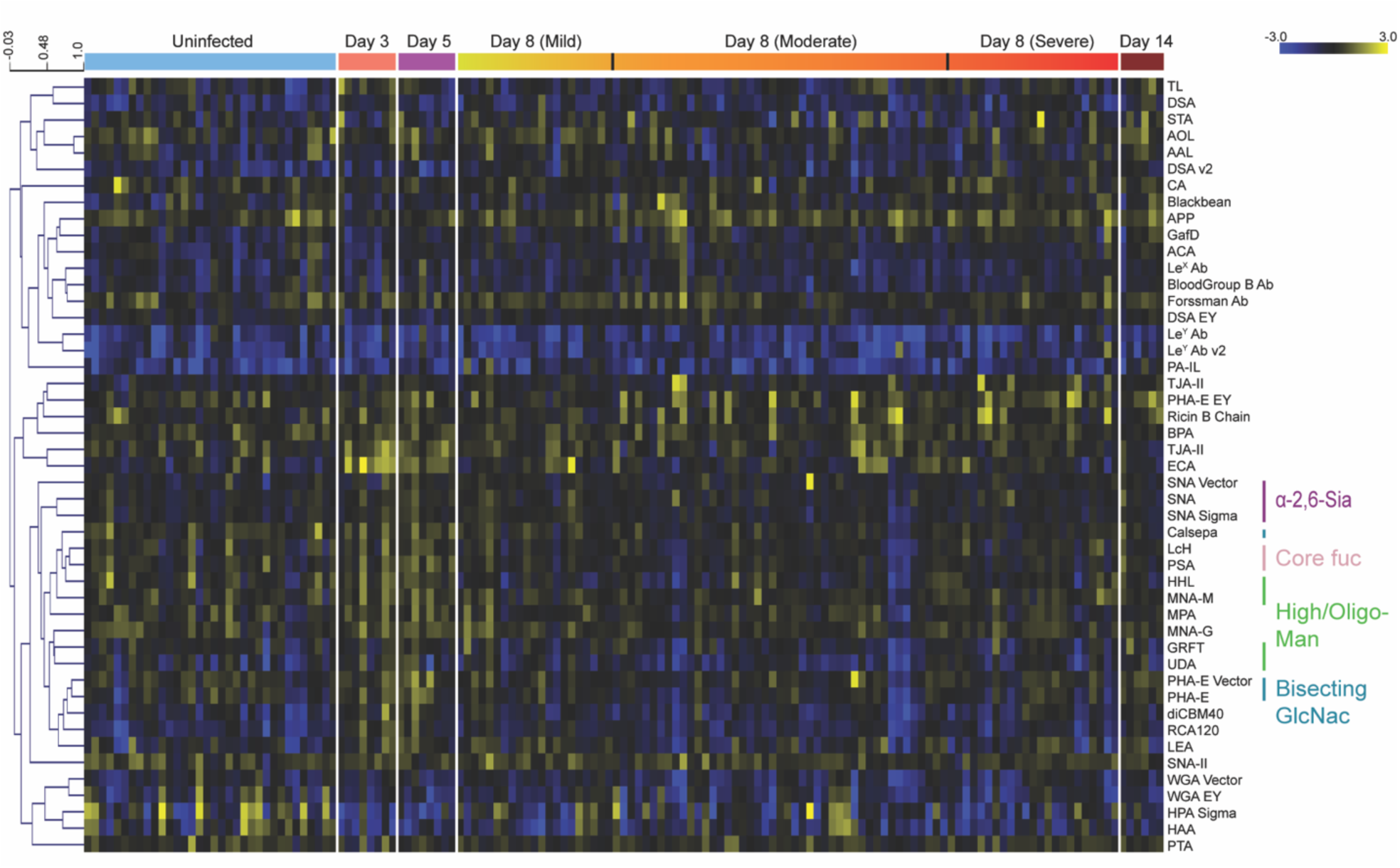
Newly weaned ferrets undergo glycan changes early in response to influenza. Median normalized log_2_ ratios (Sample (S)/Reference(R)) of newly weaned (6-7 weeks of age) ferret lung samples were ordered by days post infection. Uninfected (blue), n = 16, 2 samples per ferret; day 3 (salmon), n = 4, 2 samples per ferret; day 5 (magenta), n = 4, 2 samples per ferret; day 8, ordered by severity (mild, yellow; moderate, orange; severe, red), n = 42, 2 samples per ferret; day 14 (maroon), n = 3, 2 samples per ferret. Yellow, log_2_(S) > log_2_(R); blue, log_2_(R) > log_2_(S). Lectins binding α-2,6-sialosides (purple), high/oligo-mannose (green), bisecting GlcNac (turquoise), and core fucose (light pink) are highlighted to the right of the heatmap.

In contrast to both adults and aged ferrets, we saw no clear severity-dependent differences in the newly weaned animals at day 8 post-infection. We did observe an increase in complex N-glycan epitopes early in the course of infection, peaking by day 5 and returning to pre-infection levels by day 8 (*α*-2,6-sialic acid: SNA; core fucose: LcH, PSA; bisecting branching: PHA-E, Calsepa, **Figure 4** and **Supplementary Figure S13**). Although high/oligo-mannose trended with complex N-glycans in the time-course, the changes were not statistically significant. Overall, the lack of strong changes in the glycome is consistent with the mild nature of illness observed in young ferrets, modeling the human population.

## CONCLUSIONS

In 2009, a new pandemic H1N1 influenza A virus emerged and displayed an unusual severity pattern, impacting adults more severely than young children or the elderly. The lower impact on the elderly population, who are typically at high-risk for influenza severity, was later determined to be a result of pre-existing immunity. The ferret model of influenza has been found to mimic the differences in age-dependent illness observed in the absence of pre-existing immunity. In this model, there is a gradient of severity with young ferrets showing mild illness and aged ferrets displaying high severity. The origins of this disparity in host response is still unclear. In this work, we show dramatic age-dependent differences in the ferret glycome and host-response to influenza that overlay differences observed in severity. These include age-dependent differences in mucin-related epitopes that may explain differences in viral clearance observed between populations. In previous work, we posited that the induction of high-mannose due to influenza infection could be causative of severity through over-engagement of the innate immune system via lectins, such as MBL2. Our current study shows that high-mannose is induced in the aged population, where high levels of severity are observed, but not in newly weaned ferrets, which display a mild phenotype. This is consistent with our hypothesis and points to the need for further study to determine if the glycomic host response is a critical mediator of influenza severity.

## ASSOCIATED CONTENT

### Supporting Information

Boxplots of *α*-2,6-sialoside signal by age in uninfected ferrets; Boxplots of *α*-2,3-sialoside levels by age in uninfected ferrets; Boxplots of mucins expressions by age in uninfected ferrets; Boxplots of Lewis structures by age in uninfected ferrets; Boxplots of high/oligo-mannose levels by age in uninfected ferrets; Boxplots of Pneumonia Composite Score (PCS) as a function of weight loss quartiles in adult and aged ferrets; Aged ferrets had delayed response to influenza infection; Impact of influenza infection on α-2,6-sialosides in adult and aged ferrets; Impact of influenza infection on α-2,3-sialosides in adult and aged ferrets; Impact of influenza infection on *O*-linked glycans in aged ferrets; Dynamic changes in *O*-linked glycans are observed upon infection in aged ferrets; Comparison of severity metrics used in adult and aged ferrets; Glycan changes are observed early in the time-course study of newly weaned ferrets; Lectins used in microarrays. (PDF)

## ACKNOWLEDGMENT

The University of Georgia Institutional Animal Care and Use Committee approved all experiments under the Animal Use Protocol #2015-04-007, which were conducted in accordance with the National Research Council’s Guide for the Care and Use of Laboratory Animals, The Animal Welfare Act, and the CDC/NIH’s Biosafety in Microbiological and Biomedical Laboratories guide. This work was supported by NIAID/NIH U01 AI111598 to E. Ghedin, B. Zhang, and L. Mahal. Publication of this research was supported, in part, thanks to funding from the Canada Excellence Research Chairs Program (L. Mahal).

## Supporting Information

**Scheme 1.**
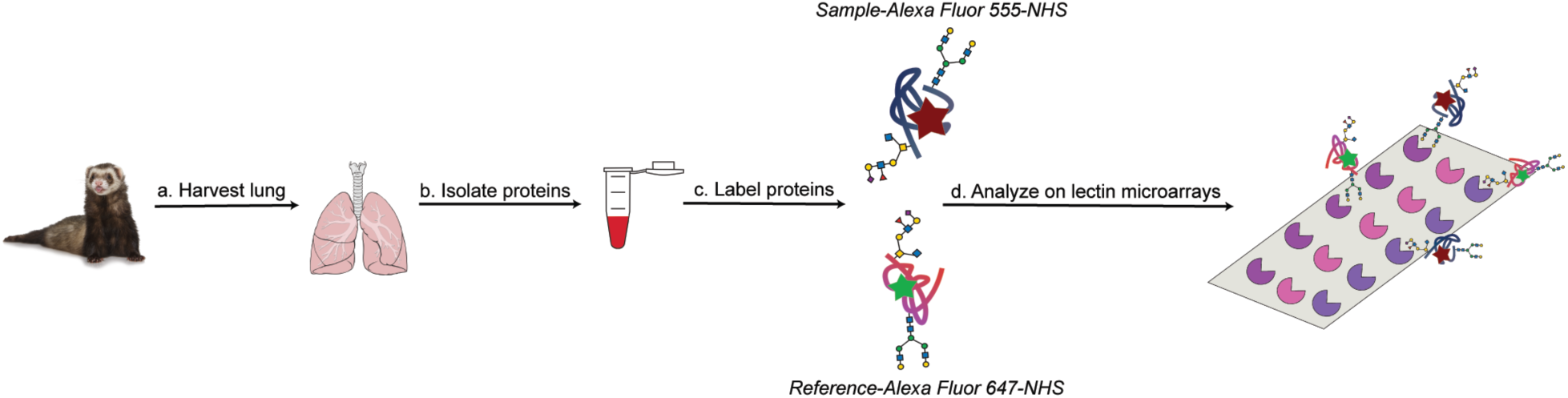
Workflow of sample preparation for dual-color lectin microarray analysis. a) Lungs were harvested from ferrets infected with H1N1pdm09 and controls. b) Glycoproteins were isolated from tissues and c) labeled with Alexa Fluor 555-NHS. A pooled reference sample was orthogonally labeled with Alexa Fluor 647-NHS. d) Equal amounts of sample and reference were mixed and analyzed on lectin microarrays.

**Scheme 2.**
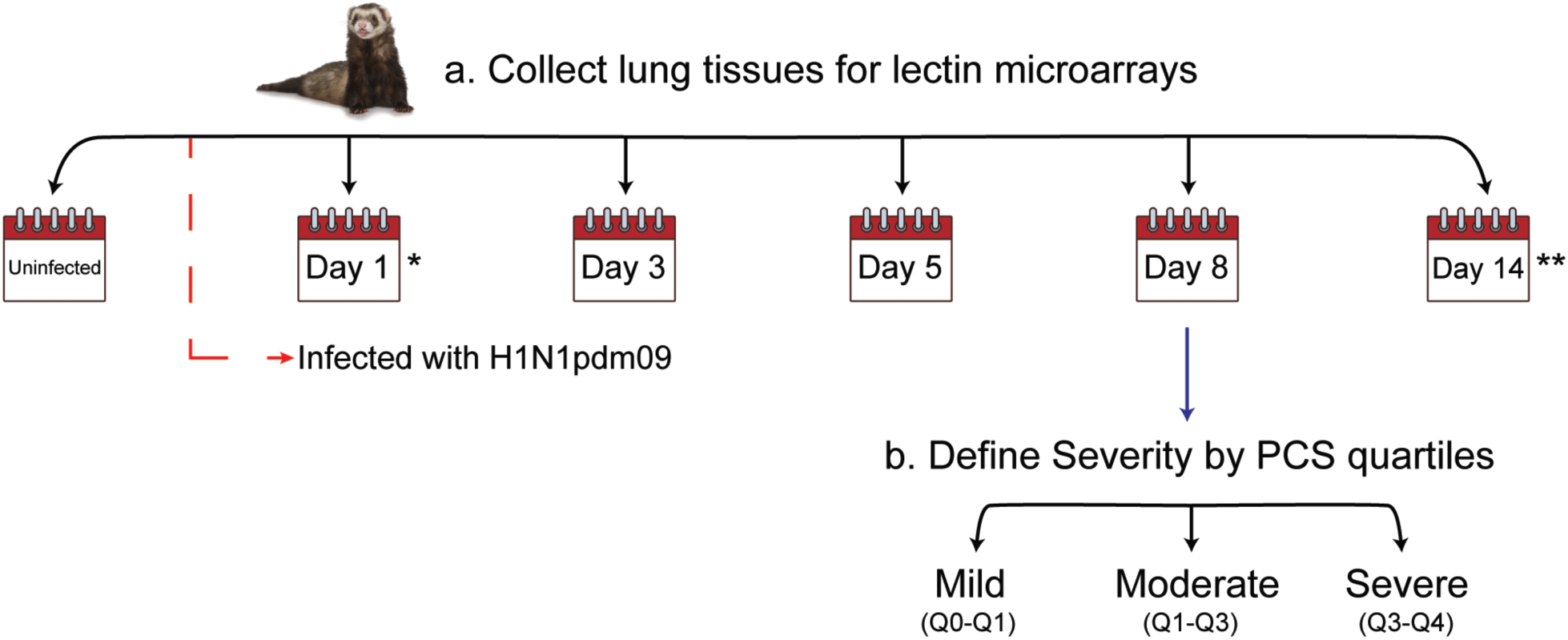
Scheme 2. Workflow of lung tissues for time-course study and severity analysis. a) Lung tissues for aged and newly weaned ferrets were collected. Lung tissues were harvested at different time points post-infection with H1N1pdm09 for lectin microarray analysis. *Day 1 time point was not collected for newly weaned ferrets. **Aged animals were sacrificed at day 8 due to weight loss >20%. b) Severity was defined by Pneumonia Composite Score (PCS) quartiles for each age group at day 8 post-infections.

**Supplementary Figure S1.**
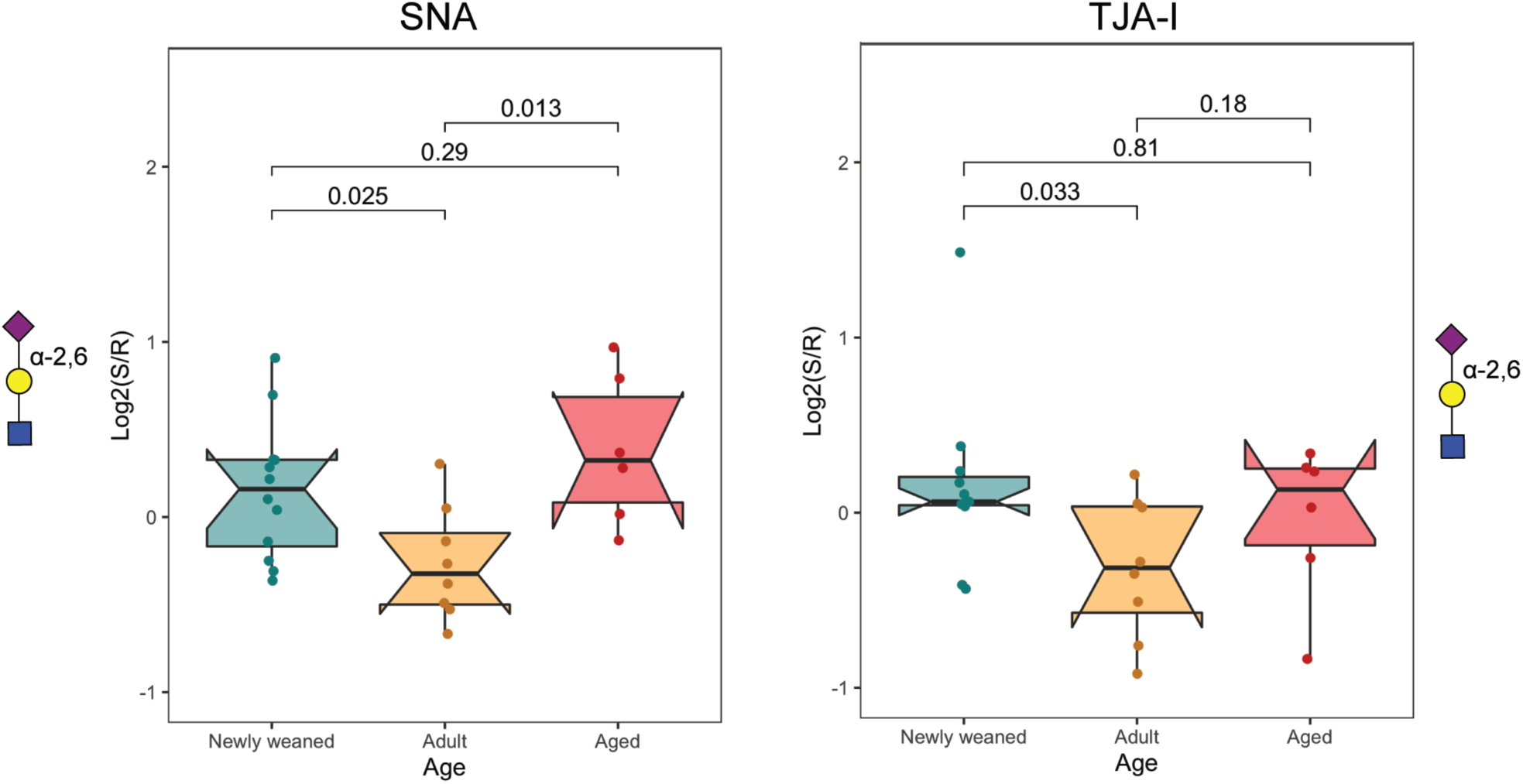
Boxplots of *α*-2,6-sialoside signal by age in uninfected ferrets. Boxplots of binding by *α*2,6-sialic acid lectins (SNA and TJA-I). Median normalized log_2_ ratios (Sample (S)/Reference(R)) are plotted by age (Newly weaned: cyan; Adult: orange; Aged: red). P-value from Wilcoxon’s t-test is shown for indicated pairs. Glycans bound by lectins are shown in the Symbolic Nomenclature for Glycomics (SNFG) at the side of the boxplots.

**Supplementary Figure S2.**
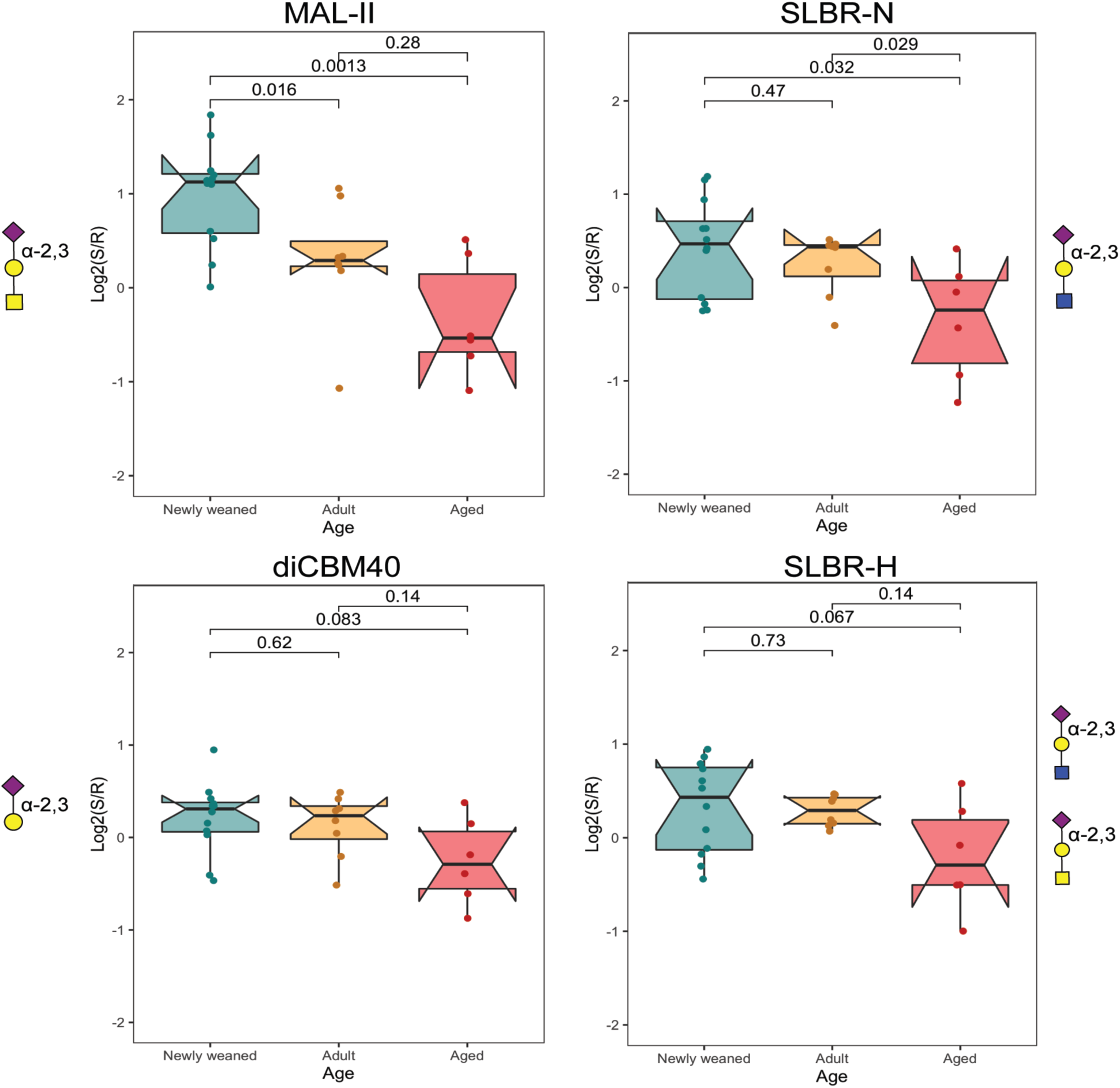
Boxplots of *α*-2,3-sialoside levels by age in uninfected ferrets. Boxplots of binding by *α*2,3-sialic acid lectins (MAL-II, SLBR-N, diCBM40, SLBR-H). Median normalized log_2_ ratios (Sample (S)/Reference(R)) are plotted by age (Newly weaned: cyan; Adult: orange; Aged: red). P-value from Wilcoxon’s t-test is shown for indicated pairs. Glycans bound by lectins are shown in SNFG at the side of the boxplots.

**Supplementary Figure S3.**
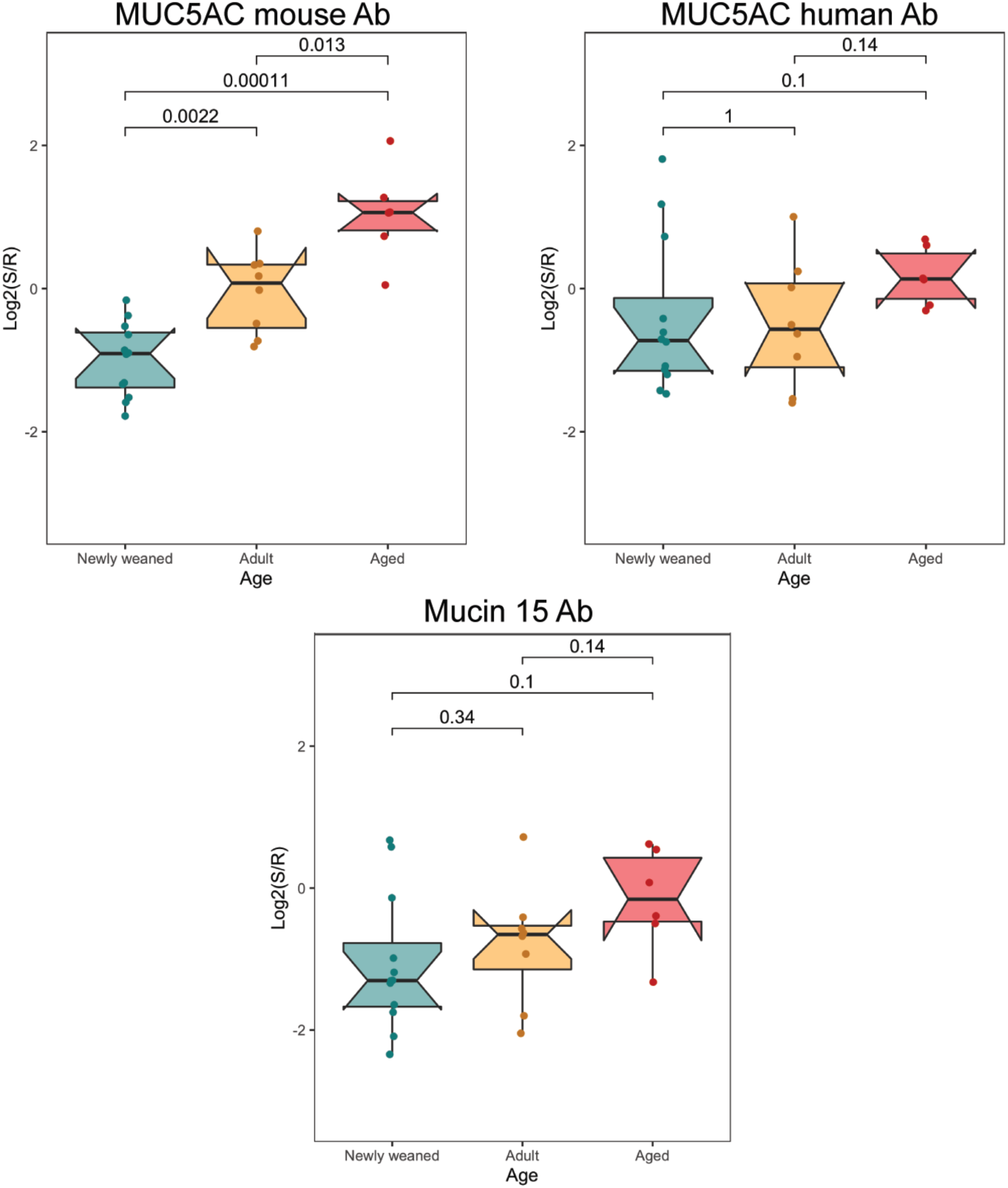
Boxplots of mucins expressions by age in uninfected ferrets. Boxplots of binding by mucin antibodies(MUC5AC mouse Ab, MUC5AC human Ab and Mucin 15 Ab). Median normalized log_2_ ratios (Sample (S)/Reference(R)) are plotted by age (Newly weaned: cyan; Adult: orange; Aged: red). P-value from Wilcoxon’s t-test is shown for indicated pairs.

**Supplementary Figure S4.**
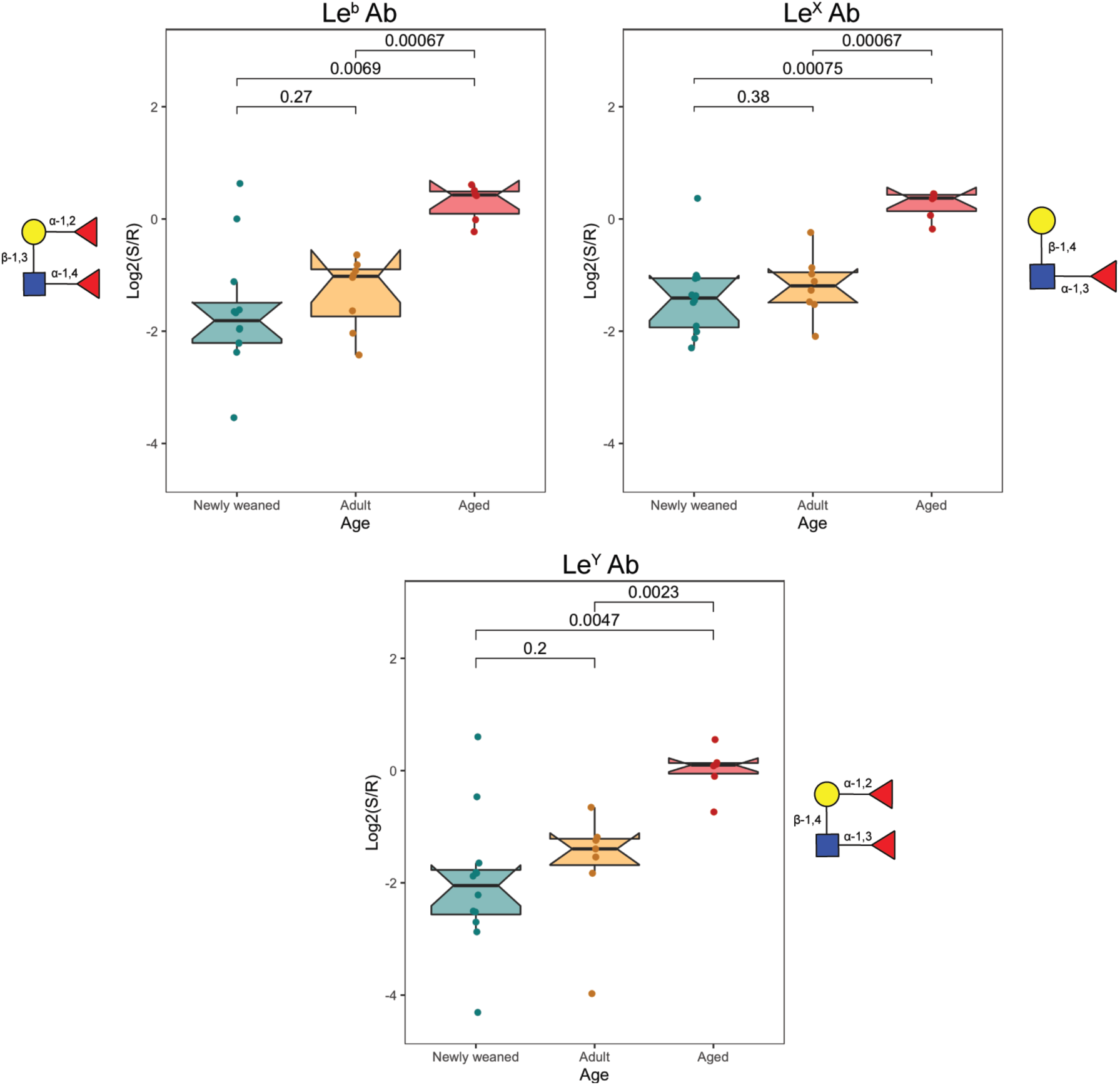
Boxplots of Lewis structures by age in uninfected ferrets. Boxplots of binding by Lewis structures antibodies(Le^b^ Ab, Le^X^ Ab and Le^Y^ Ab). Median normalized log_2_ ratios (Sample (S)/Reference(R)) are plotted by age (Newly weaned: cyan; Adult: orange; Aged: red). P-value from Wilcoxon’s t-test is shown for indicated pairs. Glycans bound by lectins are shown in SNFG at the side of the boxplots.

**Supplementary Figure S5.**
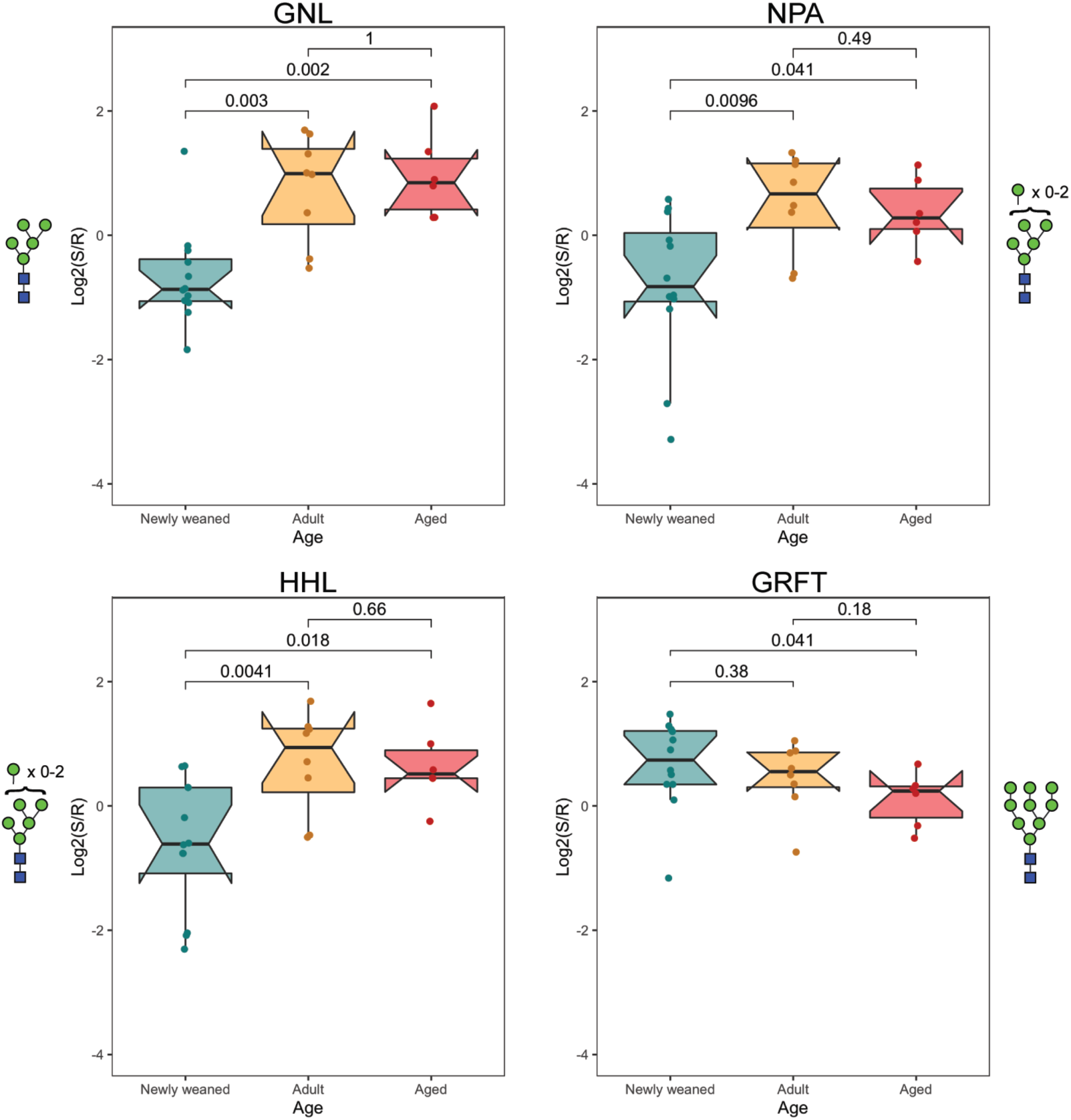
Boxplots of high/oligo-mannose levels by age in uninfected ferrets. Boxplots of binding by high/oligo-mannose lectins(GNL, NPA, HHL and GRFT). Median normalized log_2_ ratios (Sample (S)/Reference(R)) are plotted by age (Newly weaned: cyan; Adult: orange; Aged: red). P-value from Wilcoxon’s t-test is shown for indicated pairs. Glycans bound by lectins are shown in SNFG at the sides of the boxplots.

**Supplementary Figure S6.**
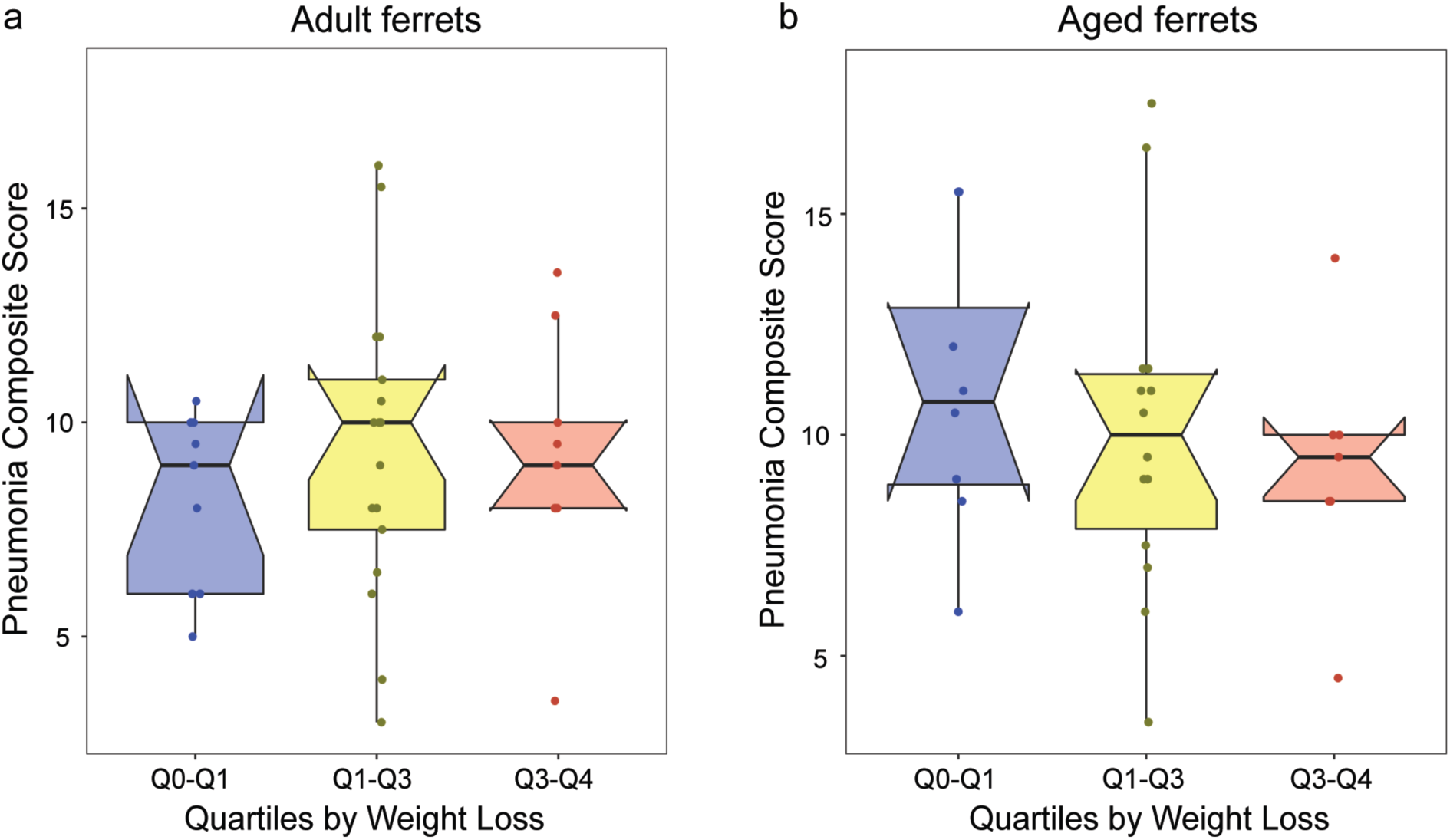
Boxplots of Pneumonia Composite Score (PCS) as a function of weight loss quartiles in adult and aged ferrets. Q0-Q1: 1^st^ quartile (blue), Q1-Q3: middle quartiles (yellow), Q3-Q4: last quartile (red) of weight loss percent. a) Boxplot analysis of PCS correlation to weight loss in adult ferrets at day 8 post-infection with H1N1pdm09. (Q0-Q1, n = 9 ferrets; Q1-Q3, n = 17; Q3-Q4, n = 9). b) Boxplot analysis of correlation in aged ferrets at day 8 post-infection (Q0-Q1, n = 8 ferrets; Q1-Q3, n = 14; Q3-Q4, n = 7).

**Supplementary Figure S7.**
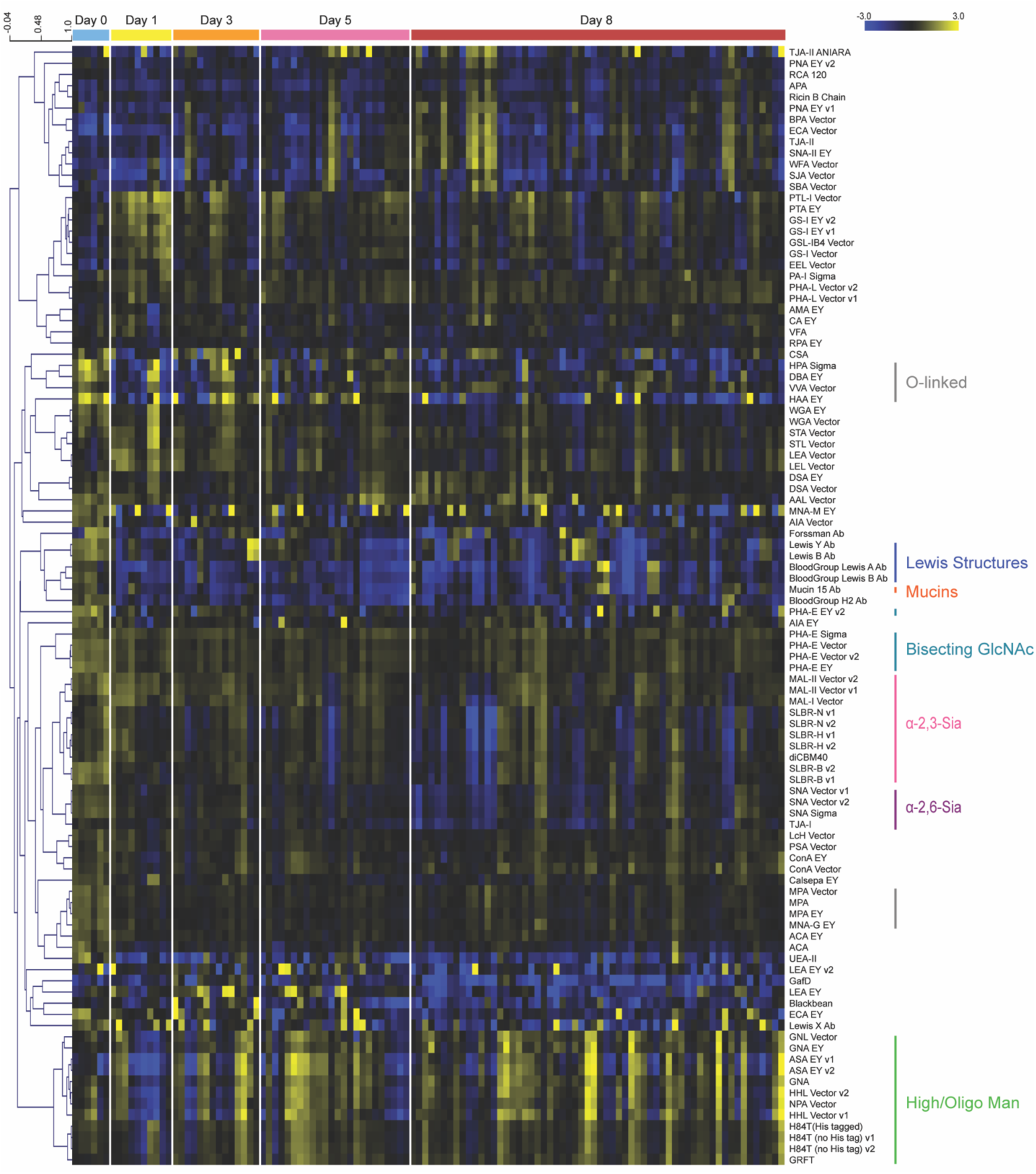
Aged ferrets had delayed response to influenza infection. Median normalized log_2_ ratios (Sample (S)/Reference(R)) of aged (>5.5 years) ferret lung samples were ordered by days post infection. Uninfected (blue), n = 3; day 1 (yellow), n = 5; day 3 (orange), n = 7; day 5 (pink), n = 12; day 8 (red), n = 29. 2 samples per ferret. Yellow, log_2_(S) > log_2_(R); blue, log_2_(R) > log_2_(S). Lectins binding α-2,3-sialosides (pink), α-2,6-sialosides (purple), high/oligo-mannose (green), bisecting GlcNAc (turquoise), *O*-linked glycans (charcoal), mucins (orange) and Lewis structures (blue) are highlighted on the right of the heatmap.

**Supplemental Figure S8.**
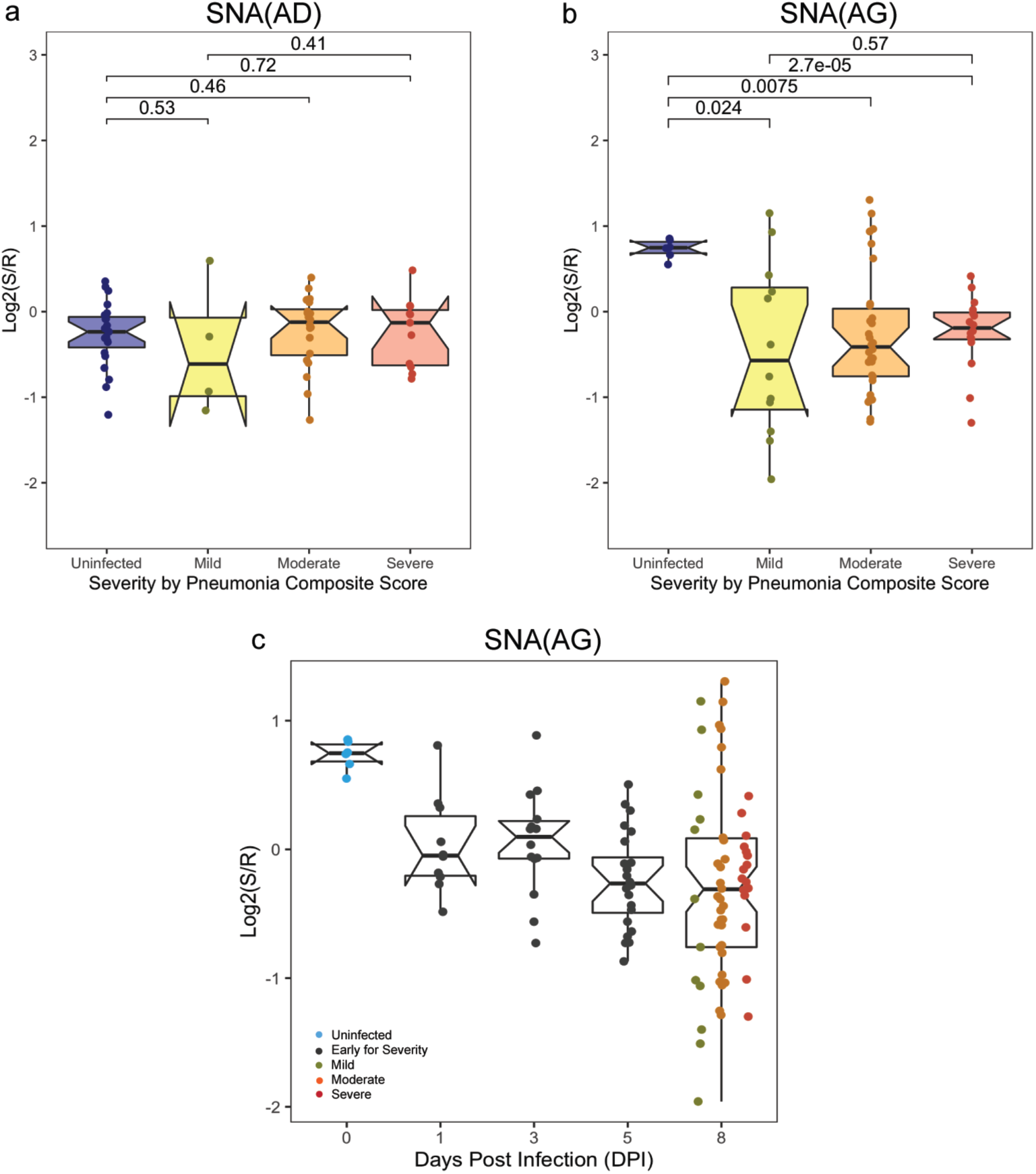
Impact of influenza infection on α-2,6-sialosides in adult and aged ferrets. a) Severity study of α-2,6-sialic acids probed by SNA at day 8 post-infection in adult ferrets (AD) is shown. Boxplots of median normalized log_2_ ratios (Sample (S)/Reference(R)) of ferret lung samples were ordered by severity defined by pneumonia composite score (PCS). Uninfected (blue): n = 4, 4 samples per ferret. Day 8 post-infection: Mild (yellow), n = 2, 2 samples per ferret; Moderate (orange), n = 11, 2 samples per ferret; Severe (red), n = 6, 2 samples per ferret. b) Severity study of α-2,6-sialic acids probed by SNA at day 8 post-infection in aged ferrets (AG) is shown. Boxplots of median normalized log_2_ ratios (Sample (S)/Reference(R)) of ferret lung samples were ordered by severity defined by PCS. Uninfected (blue): n = 3, 2 samples per ferret. Day 8 post-infection (2 samples per ferret): Mild (yellow), n = 6; Moderate (orange), n = 15; Severe (red), n = 8. P-value from Wilcoxon’s t-test is shown for indicated pairs. c) Time-course analysis of α-2,6-sialic acid levels probed by SNA following H1N1pdm09 infections (t = 0, 1, 3, 5 and 8 days) in aged animals (AG). Boxplot of median normalized log_2_ ratios (Sample (S)/Reference(R)) is shown. Disease severity is indicated by color (Uninfected: blue; Early for Severity: black; Mild: dark yellow; Moderate: orange, Severe: red).

**Supplemental Figure S9.**
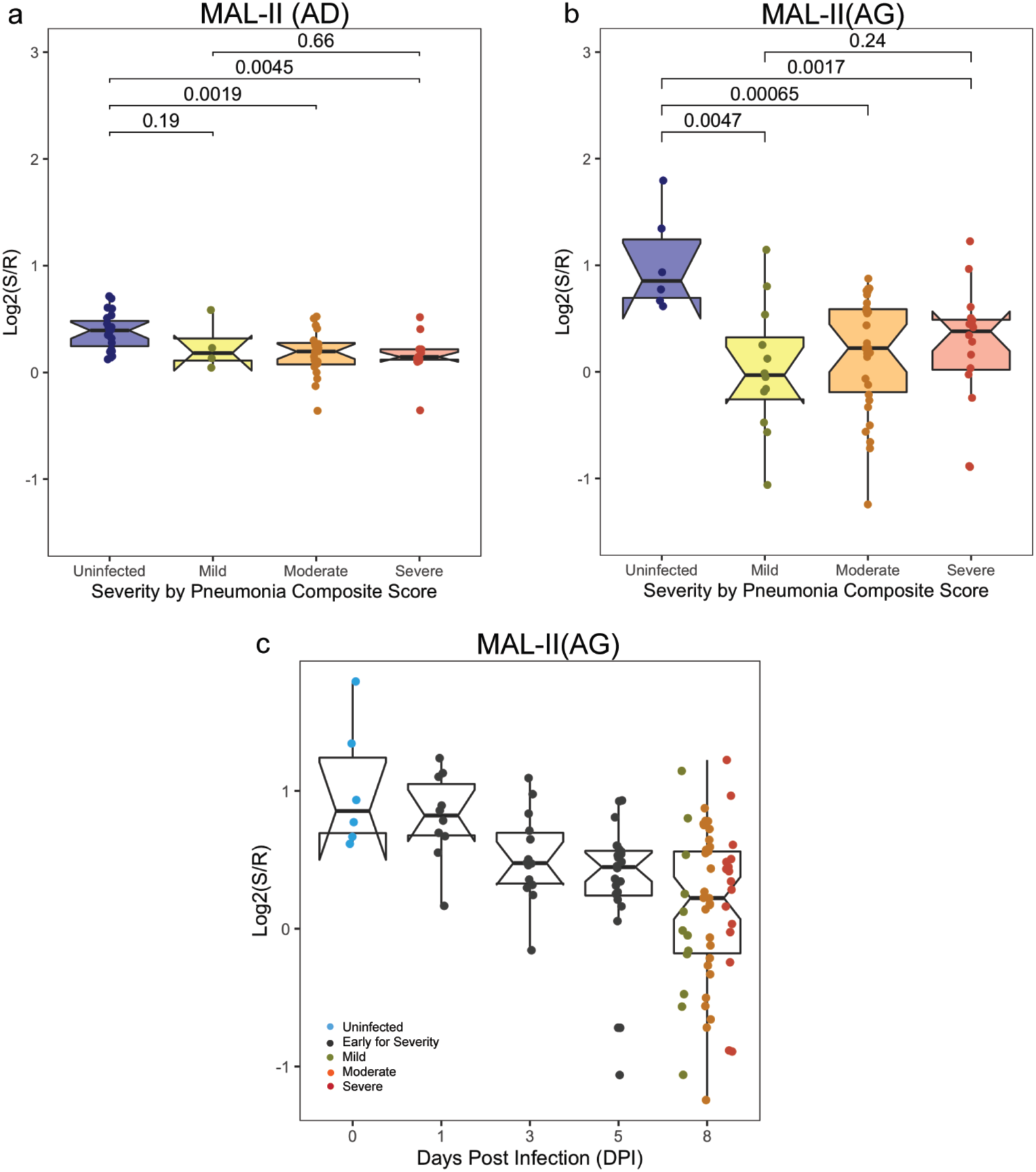
Impact of influenza infection on α-2,3-sialosides in adult and aged ferrets. a) Severity study of α-2,3-sialic acids probed by MAL-II at day 8 post-infection in adult ferrets (AD) is shown. Boxplots of median normalized log_2_ ratios (Sample (S)/Reference(R)) of ferret lung samples were ordered by severity defined by pneumonia composite score (PCS). Uninfected (blue): n = 4, 4 samples per ferret. Day 8 post-infection: Mild (yellow), n = 2, 2 samples per ferret; Moderate (orange), n = 11, 2 samples per ferret; Severe (red), n = 6, 2 samples per ferret. b) Severity study of α-2,3-sialic acids probed by MAL-II at day 8 post-infection in aged ferrets (AG) is shown. Boxplots of median normalized log_2_ ratios (Sample (S)/Reference(R)) of ferret lung samples were ordered by severity defined by PCS. Uninfected (blue): n = 3, 2 samples per ferret. Day 8 post-infection: Mild (yellow), n = 6, 2 samples per ferret; Moderate (orange), n = 15, 2 samples per ferret; Severe (red), n = 8, 2 samples per ferret. P-value from Wilcoxon’s t-test is shown for indicated pairs. c) Time-course analysis of α-2,3-sialic acid levels probed by MAL-II following H1N1pdm09 infections (t = 0, 1, 3, 5 and 8 days) in aged animals (AG). Boxplot of median normalized log_2_ ratios (Sample (S)/Reference(R)) is shown. Disease severity is indicated by color (Uninfected: blue; Early for Severity: black; Mild: dark yellow; Moderate: orange, Severe: red).

**Supplemental Figure S10.**
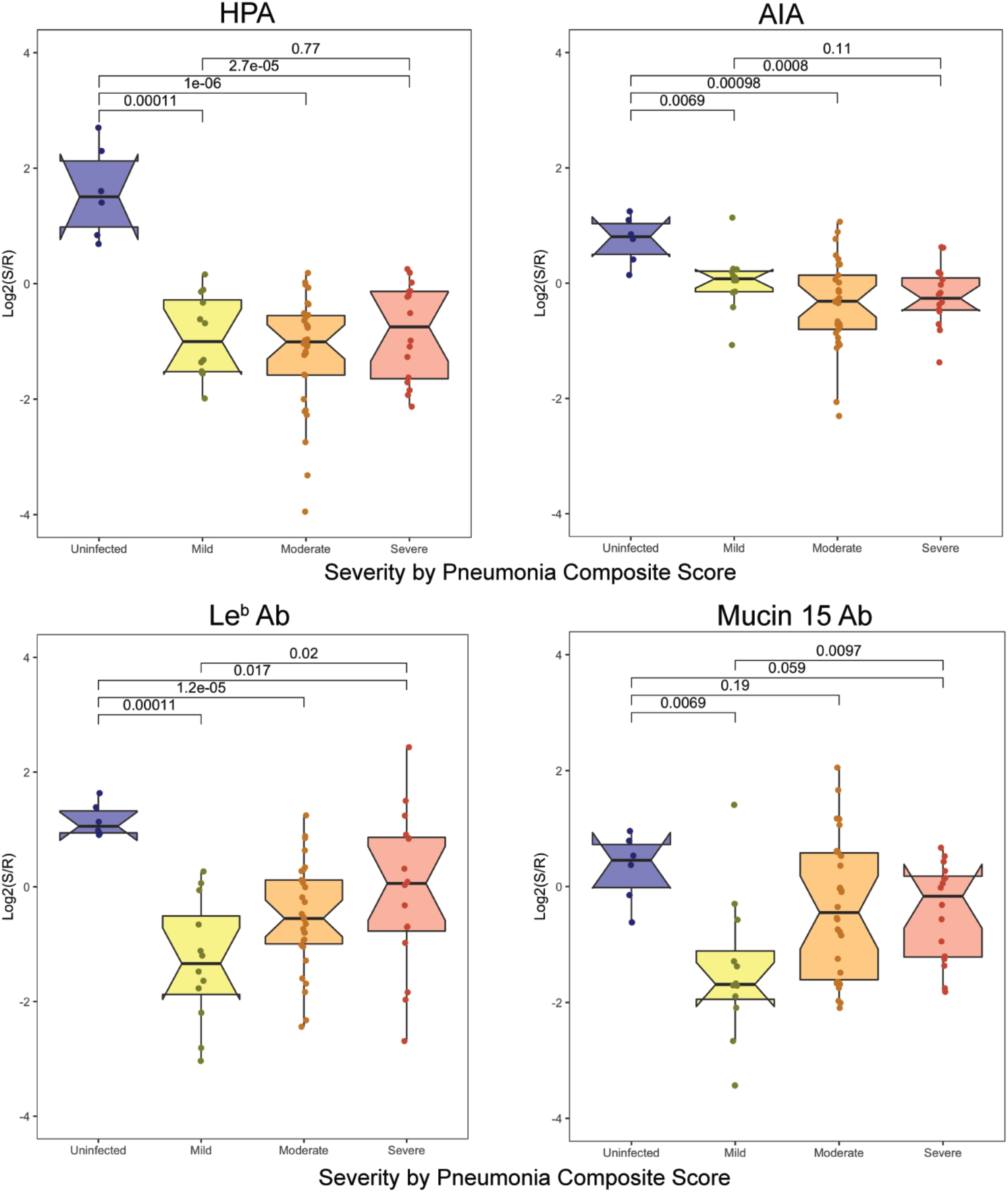
Impact of influenza infection on *O*-linked glycans in aged ferrets. Severity study of *O*-linked glycans probed by HPA and AIA, Lewis structures by Le^b^ Ab, and mucins by Mucin 15 Ab at day 8 post-infection in aged ferrets is shown. Boxplots of median normalized log_2_ ratios (Sample (S)/Reference(R)) of ferret lung samples were ordered by severity defined by PCS. Uninfected (blue): n = 3, 2 samples per ferret. Day 8 infected: Mild (yellow), n = 6, 2 samples per ferret; Moderate (orange), n = 15, 2 samples per ferret; Severe (red), n = 8, 2 samples per ferret. P-value from Wilcoxon’s t-test is shown for indicated pairs.

**Supplementary Figure S11.**
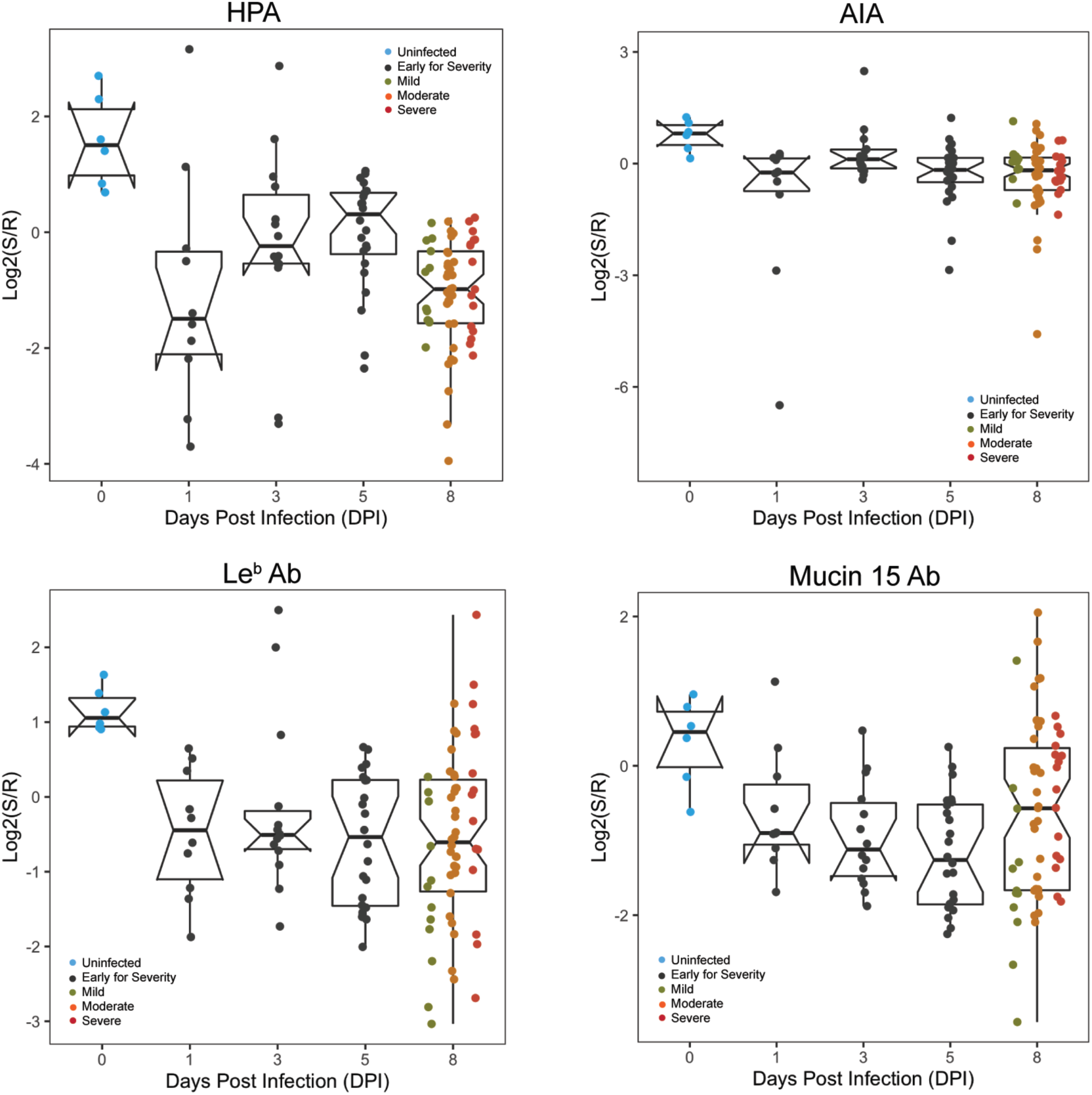
Dynamic changes in *O*-linked glycans are observed upon infection in aged ferrets. Boxplot analysis of *O*-linked glycans probed by HPA and AIA, Lewis structures probed by Le^b^ Ab, and mucins probed by Mucin 15 Ab following H1N1pdm09 infections (t = 0, 1, 3, 5 and 8 days). Boxplot of median normalized log_2_ ratios (Sample (S)/Reference(R)) is shown. Disease severity is indicated by color (Uninfected: blue; Early for Severity: black; Mild: dark yellow; Moderate: orange, Severe: red).

**Supplementary Figure S12.**
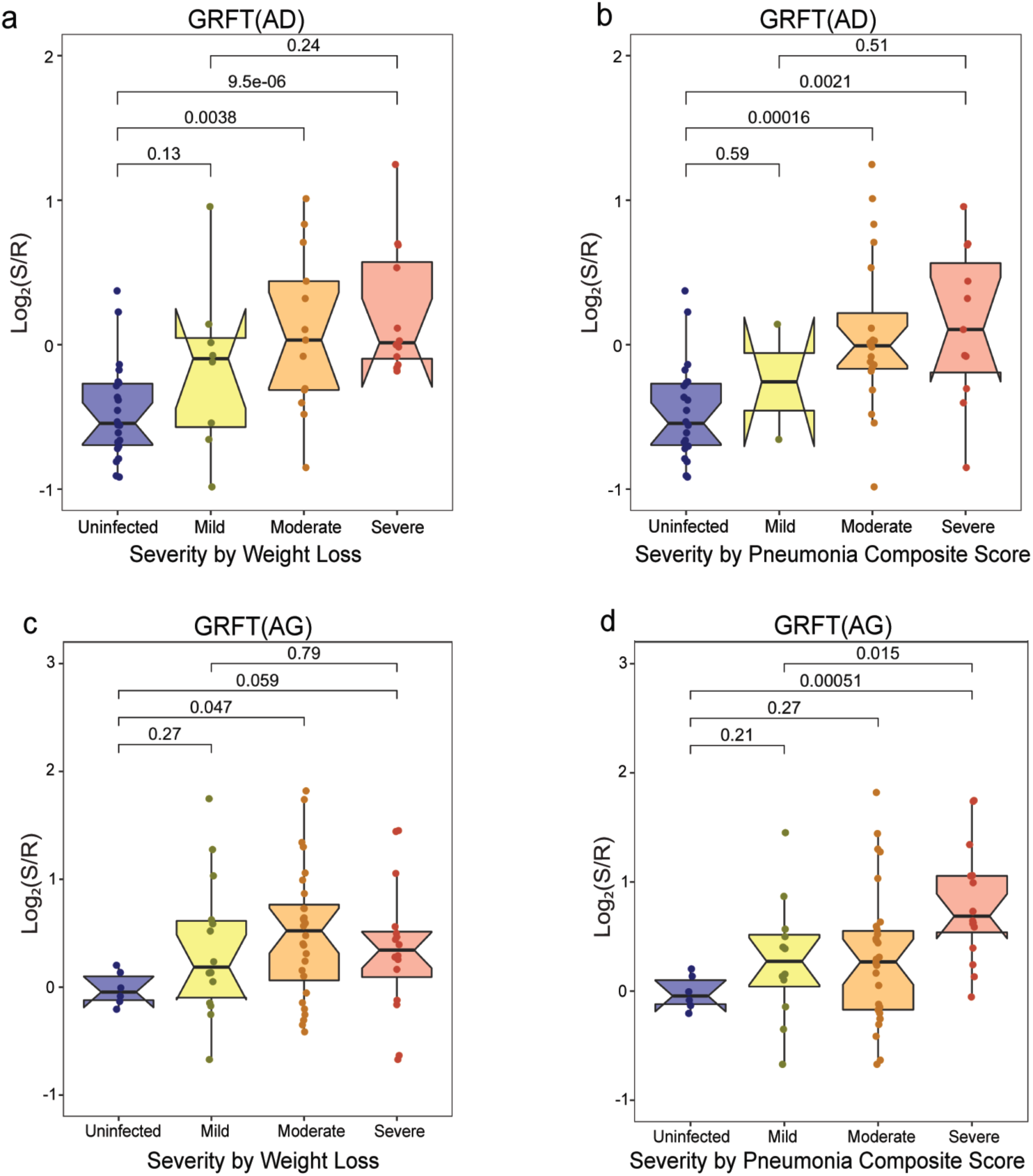
Comparison of severity metrics used in adult and aged ferrets. a) Boxplot analysis of lectin binding by GRFT (high-mannose) as a function of severity defined by weight loss in adult ferrets (AD). Uninfected (blue): n = 4, 4 samples per ferret. Day 8 post-infection: Mild (yellow), n = 5, 2 samples per ferret; Moderate (orange), n = 7, 2 samples per ferret; Severe (red), n = 7, 2 samples per ferret. b) Boxplot analysis of lectin binding by GRFT (high-mannose) as a function of severity defined by pneumonia composite score (PCS) in adult ferrets (AD). Uninfected (blue): n = 4, 4 samples per ferret. Day 8 post-infection: Mild (yellow), n = 2, 2 samples per ferret; Moderate (orange), n = 11, 2 samples per ferret; Severe (red), n = 6, 2 samples per ferret. c) Boxplot analysis of lectin binding by GRFT (high-mannose) as a function of severity defined by weight loss in aged ferrets (AG). Uninfected (blue): n = 3, 2 samples per ferret. Day 8 post-infection: Mild (yellow), n = 7, samples per ferret; Moderate (orange), n = 14, 2 samples per ferret; Severe (red), n = 8, 2 samples per ferret. d) Boxplot analysis of lectin binding by GRFT (high-mannose) as a function of severity defined by PCS in aged ferrets (AG). Uninfected (blue): n = 3, 2 samples per ferret. Day 8 post-infection: Mild (yellow), n = 6, 2 samples per ferret; Moderate (orange), n = 15, 2 samples per ferret; Severe (red), n = 8, 2 samples per ferret.

**Supplementary Figure S13.**
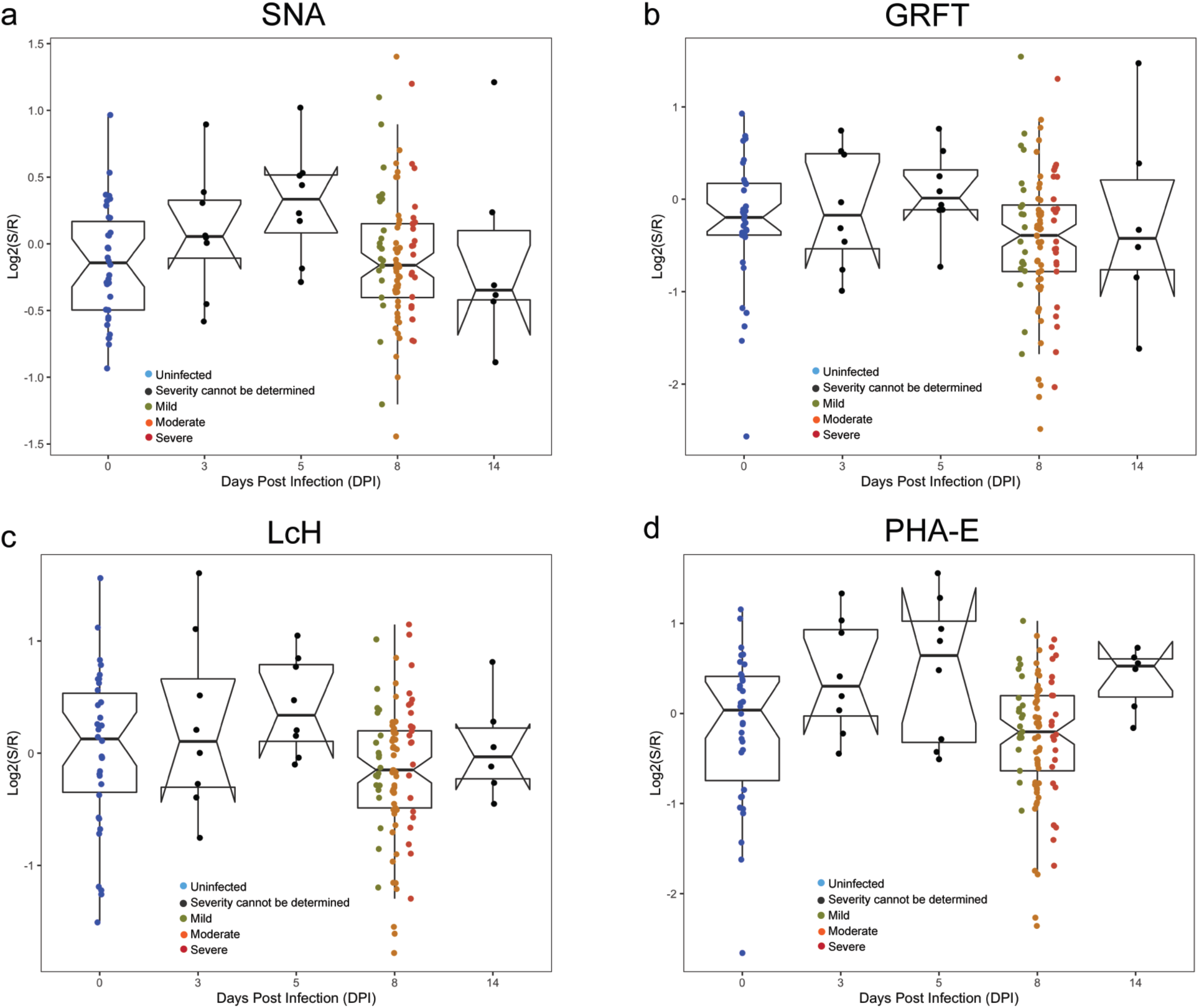
Glycan changes are observed early in the time-course study of newly weaned ferrets. a) Time-course analysis of α-2,6-sialic acid probed by SNA following H1N1pdm09 infections (t = 0, 3, 5, 8 and 14 days). b) Time-course analysis of high/oligo-mannose probed by GRFT. c) Time-course analysis of core fucose probed by LcH. d) Time-course analysis of bisecting branching probed by PHA-E. Boxplot of median normalized log_2_ ratios (Sample (S)/Reference(R)) is shown. Disease severity is indicated by color (Uninfected: blue; Severity cannot be determined: black; Mild: dark yellow; Moderate: orange, Severe: red).

**Supplemental Table 1.** Lectins used in microarrays.

| **Lectin** | **Species/Origin** | **Print Conc.**<br>**(μg/mL)** | **Rough**<br>**Specificity**<br>**/Inhibitory**<br>**monosaccharide** | **Vendor/Source** |
| --- | --- | --- | --- | --- |
| AAL <sup>a,b,c</sup> | *Aleuria aurantia* | 1000 | Fucose | Vector |
| ACA <sup>a,b,c</sup> | *Amaranthus Caudatus* | 1000 | Gal-β1,3-GalNAc | Vector |
| AIA <sup>a,b,c</sup> | *Artocarpus integrifolia* | 500 | β1,3-GalNAc | Vector/EY |
| AMA <sup>a,b,c</sup> | *Allium moly* | 500 | Oligo mannose | EY |
| Anti-B.G.B [CLCP-19B] <sup>a</sup> | MAb mouse IgM [CLCP-19B] | undiluted | Blood group B antigen | Abcam |
| Anti-B.G.H2 <sup>a,b,c</sup> | MAb mouse IgM [A46-B/B10] | undiluted | Blood group H2 antigen | Santa Cruz Biotechnology |
| Anti-CD 15[MY-1] <sup>a</sup> | Mab mouse IgM [MY-1] | undiluted | Lewis X | Abcam |
| Anti-B.G. Lewis A [7LE] <sup>b,c</sup> | MAb mouse IgG1 [7LE] | undiluted | Lewis A | Abcam |
| Anti-B.G. Lewis B [2-25LE] <sup>b,c</sup> | MAb mouse IgM [2-25LE] | undiluted | Lewis B | Abcam |
| Anti-B.G.Lewis Y <sup>b,c</sup> | MAb mouse IgM | undiluted | Lewis Y | Santa Cruz Biotechnology |
| Anti-Forssman [117C9] <sup>a,b,c</sup> | MAb Rat IgM [117C9] | undiluted | Forssman Antigen | Abcam |
| Anti-Lewis B [T218] <sup>a,b,c</sup> | IgM [T218] | undiluted | Lewis B | Sigma |
| Anti-Lewis X [P12] <sup>a,b,c</sup> | MAb mouse IgM [P12] | undiluted | Lewis X | Abcam |
| Anti-Lewis Y [F3] <sup>a,b,c</sup> | MAb mouse IgM [F3] | undiluted | Lewis Y | Abcam |
| Anti-MUC5AC human [CLH2] <sup>b</sup> | Mab mouse IgG1 [CLH2] | undiluted | human MUC5AC | Sigma |
| Anti-MUC5AC mouse <sup>b</sup> | Goat polyclonal to mouse MUC5AC | undiluted | mouse MUC5AC | LSBio |
| Anti-Mucin 15 [H-5] <sup>b,c</sup> | Mab mouse IgG1 [H-5] | undiluted | Mucin 15 | Santa Cruz Biotechnology |
| AOL <sup>a,b,c</sup> | *Aspergillus oryzae* | 1000 | Fucose | TCI America |
| APA <sup>a,b,c</sup> | *Abrus precatorius* | 500 | Gal-β1,3-GalNAc / Lac | EY |
| APP <sup>a</sup> | *Aegopodium podagraria* | 1000 | GalNAc / Lac / Gal | EY |
| ASA <sup>a,b,c</sup> | *Allium sativum* | 1000 | Mannose | EY |
| BDA <sup>a</sup> | *Bryonia dioica* | 1000 | GalNAc / Lac | EY |
| Blackbean <sup>a,b,c</sup> | *Blackbean crude* | 1000 | GalNAc | EY |
| BPA <sup>a,b,c</sup> | *Bauhinia purpurea* | 500 | β-Gal / β-GalNAc | Vector |
| CA <sup>a,b,c</sup> | *Colchicum autumnale* | 1200 | Bi-antennary N-linked glycans | EY |
| CAA <sup>a</sup> | *Caragana arborescens* | 1000 | Bi-antennary N-linked glycans | EY |
| Calsepa <sup>b,c</sup> | *Calystegia sepium* | 1000 | Bisecting N-linked glycans | EY |
| CCA <sup>a</sup> | *Cancer antennarius* | 1000 | 9-O-Acetyl sialylation / 4-O-Acetyl sialylation | EY |
| Cholera Toxin <sup>b</sup> | *Vibrio cholerae* | 1000 | GM1 ganglioside | Sigma |
| Con A <sup>b,c</sup> | *Canavalia ensiformis* | 1000 | Tri-mannose core | EY/Vector |
| CSA <sup>a,b,c</sup> | *Cystisus scoparius* | 1000 | Terminal GalNAc | EY |
| DBA <sup>a,b,c</sup> | *Dolichos Biflorus* | 1000 | GalNAc | Vector |
| diCBM40 <sup>a,b,c</sup> | engineered NanI from *Clostridium perfringens* | 1000 | $\alpha$ Sialylation | Generated in house |
| DSA <sup>a,b,c</sup> | *Datura stramonium* | 500 | LacNAc | EY/Vector |
| ECA <sup>a,b,c</sup> | *Erythrina cristagalli* | 1000 | LacNAc | Vector |
| EEL/EEA <sup>a,b,c</sup> | *Eunonymus europaeus* | 1000 | Blood Group B | Vector/EY |
| GafD <sup>a,b,c</sup> | recombinant GafD from *Escherichia coli* | 1000 | GlcNAc | Generated in house |
| GNA/GNL <sup>a,b,c</sup> | *Galanthus nivalis* | 1500 | Oligo mannose | Vector/EY |
| GHA <sup>a</sup> | *Glechoma hederacea* | 500 | GalNAc | EY |
| GS-I <sup>a,b,c</sup> | *Griffonia simplicifolia-I* | 1000 | α-Gal / Lac | Vector/EY |
| GS-II <sup>a,b,c</sup> | *Griffonia simplicifolia-II* | 1000 | GlcNAc | Vector |
| GS-IB4 <sup>a,b,c</sup> | *Griffonia simplicifolia-I, isolectin B4* | 2000 | Gal | Vector |
| H84T <sup>b,c</sup> | *Banana lectin* | 1000 | High mannose | Gift from Dr. David Markovitz |
| HAA <sup>a,b,c</sup> | *Homarus americanus* | 1000 | Terminal GalNAc | EY |
| HHL <sup>a,b,c</sup> | *Hippeastrum Hybrid* | 1500 | Oligo/High mannose | Vector |
| HPA <sup>a,b,c</sup> | *Helix pomatia* | 1000 | Blood Group A | Sigma/EY |
| IRA <sup>a</sup> | *Iris Hybrid* | 1000 | GalNAc / Lac | EY |
| LBA <sup>a</sup> | *Phaseolus lunatus* | 1000 | Blood Group A | EY |
| LAA <sup>b</sup> | *Laburnum alpinum* | 900 | GlcNAc | EY |
| LcH <sup>a,b,c</sup> | *Lens Culinaris* | 1000 | Core Fucose | Vector |
| LEA/LEL <sup>a,b,c</sup> | *Lycopersicon esculentum* | 1000 | GlcNAc | Vector/EY |
| LFA <sup>a</sup> | *Limax flavus* | 500 | $\alpha$ Sialylation | EY |
| Lotus <sup>a,b,c</sup> | *Lotus tetragonolobus* | 1000 | Fucose | Vector |
| MAA <sup>a</sup> | *Maackia amurensis* | 500 | Sialylation/Sulfation | EY |
| MAL-I <sup>a,b,c</sup> | *Maackia amurensis-I* | 2000 | Sialylation/Sulfation | Vector |
| MAL-II <sup>a,b,c</sup> | *Maackia amurensis-II* | 2000 | Sialylation/Sulfation | Vector |
| MNA-G <sup>a,b,c</sup> | *Morus nigra Morniga G* | 1000 | GalNAc | EY |
| MNA-M <sup>a,b,c</sup> | *Morus nigra Morniga M* | 1000 | Oligo mannose / Gal | EY |
| MPA/MPL <sup>a,b,c</sup> | *Maclura pomifera* | 1000 | β1,3-GalNAc | Vector |
| newBR6 <sup>b</sup> | unknown (from unpublished work) | 480 | under investigation | Gift from Dr. Barbara Bensing |
| NPA <sup>a,b,c</sup> | *Narcissus pseudonarcissus* | 1000 | Oligo mannose | Vector |
| PA-I <sup>a,b,c</sup> | *Pseudomonas aeruginosa* | 1000 | Gal | Sigma |
| PA-IL <sup>a</sup> | *bacteria* | 1000 | GalNAc | Generated in house |
| PHA-E <sup>a,b,c</sup> | *Phaseolus vulgaris Erythroagglutinin* | 1000 | Bisecting GlcNAc | Vector/EY/Sigma |
| PHA-L <sup>a,b,c</sup> | *Phaseolus vulgaris Leukoagglutinin* | 1000 | β1,6 Branching N-Link glycans | Vector/EY |
| PMA <sup>a</sup> | *Polygonatum multiflorum* | 500 | Oligo mannose | EY |
| PNA <sup>a,b,c</sup> | *Arachis hyogaea* | 1000 | Gal-β1,3-GalNAc | Vector/EY |
| PSA <sup>a,b,c</sup> | *Pisum sativum* | 1000 | Core Fucose | Vector |
| PSL <sup>a</sup> | *Polyporus squamosus* | 1000 | α2,6 sialylation | EY |
| PTA <sup>a,b,c</sup> | *Psophocarpus tetragonolobus* | 500 | Blood Groups | EY |
| PTL-I <sup>a,b,c</sup> | *Psophocarpus tetragonolobus-I* | 1500 | Blood Group A | Vector |
| PTL-II <sup>a,b,c</sup> | *Psophocarpus tetragonolobus-II* | 1000 | α2 Fucose | Vector |
| RCA120 <sup>a,b,c</sup> | *Ricinus Communis Agglutinin I* | 1000 | Gal / Lac | Vector |
| rCVN <sup>a</sup> | *recombinant Cyanovirin* | 1000 | High mannose | Gift from Dr. Barry O'Keefe |
| rGRFT <sup>a,b,c</sup> | *recombinant Griffithsin* | 1000 | High mannose | Gift from Dr. Barry O'Keefe |
| Ricin B Chain <sup>a,b,c</sup> | *Ricinus communis* | 1000 | Gal | Vector |
| RPA <sup>a,b,c</sup> | *Robinia pseudoacacia* | 500 | Complex N-link glycans | EY |
| rSVN <sup>a,b</sup> | *recombinant Scytovirin* | 1000 | High mannose | Gift from Dr. Barry O'Keefe |
| SBA <sup>a,b,c</sup> | *Glycine max* | 1000 | LacdiNAc | Vector |
| SJA <sup>a,b,c</sup> | *Sophora japonica* | 1000 | LacdiNAc | Vector |
| SK1 <sup>b</sup> | *Streptococcus sanguinis SK1* | 1800 | α2,3 sialylation | Gift from Dr. Barbara Bensing |
| SK678 <sup>b</sup> | *Streptococcus sanguinis SK678* | 450 | α2,3 sialylation | Gift from Dr. Barbara Bensing |
| SLBR-B <sup>b,c</sup> | *Streptococcus gordonii M99* | 1000 | α2,3 sialylation | Gift from Dr. Barbara Bensing |
| SLBR-H <sup>b,c</sup> | *Streptococcus gordonii DL1* | 2000 | α2,3 sialylation | Gift from Dr. Barbara Bensing |
| SLBR-N <sup>b,c</sup> | *Streptococcus gordonii UB10712* | 1000 | α2,3 sialylation | Gift from Dr. Barbara Bensing |
| SNA <sup>a,b,c</sup> | *Sambucus nigra* | 500/1000 | α2,6 sialylation | Vector/Sigma |
| SNA-II <sup>a,b,c</sup> | *Sambucus nigra-II* | 1000 | α2 Fucose /oligo mannose | EY |
| STA/STL <sup>a,b,c</sup> | *Solanus tuberosum* | 500 | GlcNAc | Vector |
| TJA-I <sup>a,b,c</sup> | *Trichosanthes japonica-I* | 1000 | α2,6 sialylation | TCI |
| TJA-II <sup>a,b,c</sup> | *Trichosanthes japonica-II* | 1000 | α2 Fucose | NorthStar Bioproducts/Ani ara Diagnostica |
| TL <sup>a,b,c</sup> | *Tulipa sp.* | 700 | GlcNAc | EY |
| UDA <sup>a,b</sup> | *Urtica dioica* | 1000 | GlcNAc / Oligo mannose | EY |
| UEA-I <sup>a,b,c</sup> | *Ulex europaeus-I* | 1000 | α2 Fucose | Vector |
| UEA-II <sup>a,b,c</sup> | *Ulex europaeus-II* | 2000 | GlcNAc | Vector |
| VFA <sup>a,b,c</sup> | *Vicia faba* | 1000 | GlcNAc | EY |
| VVA <sup>a,b,c</sup> | *Vicia villosa* | 1000 | Terminal GalNAc | Vector/EY |
| VVA(man) <sup>a,b,c</sup> | *Vicia villosa* | 500 | Mannose | Vector/EY |
| WFA <sup>a,b,c</sup> | *Wisteria floribunda* | 1000 | GalNAc-β1,4 | Vector |
| WGA <sup>a,b,c</sup> | *Triticum vulgare* | 1000 | GlcNAc | Vector/EY |
| X408 <sup>b</sup> | unknown (from unpublished work) | 1200 | under investigation | Gift from Dr. Barbara Bensing |

<sup>a</sup> : lectins printed in uninfected experiment
<sup>b</sup> : lectins printed in newly weaned experiment
<sup>c</sup> : lectins printed in aged experiment

